# Evaluating Deep Learning Based Structure Prediction Methods on Antibody-Antigen Complexes

**DOI:** 10.1101/2025.07.11.662141

**Authors:** Samuel Fromm, Marko Ludaic, Arne Elofsson

## Abstract

**Motivation:** AlphaFold2 significantly improved the prediction of protein complex structures. However, its accuracy is lower for interactions without coevolutionary signals, such as host-pathogen and antibody-antigen interactions. Two strategies have been developed to address this limitation: massive sampling and replacing the evoformer with the pairformer, which does not rely on coevolution, as introduced in AlphaFold3, thereby enabling more structural reasoning by the network.

**Results:** In this study, we benchmark structure prediction methods on unseen antibody-antigen complexes. We found that increased sampling improves the chances of generating a correct protein model, roughly in a log-linear manner. However, the internal quality estimates by AlphaFold often cannot identify the best predicted structures for each target, resulting in a significant loss of performance for the top-ranked protein model compared with the best model. For all methods, a significant challenge remains the identification of the best model. We also show that AlphaFold3 outperforms AlphaFold2, Boltz-1, and Chai-1. Furthermore, AlphaFold3 performance declines significantly for complexes that lack structural similarity to the training set, indicating that it has to some extent learned to detect remote structural similarities.

**Availability and implementation:** All code is available from github.com/samuelfromm/abag-benchmark-set/ and all data from `DOI:10.5281/zenodo.17978681`. The latter repository also contains the code.

**Supplementary information:** Supplementary information is available online.

## Introduction

Antibody-antigen interactions play a crucial role in immunology and various biotechnological applications. Furthermore, the use of therapeutic antibodies has increased over the past few decades (1). Therefore, accurate prediction of these interactions is essential. It can accelerate discovery by identifying potential immunotherapy candidates and enhancing our ability to design targeted treatments for various diseases, including cancers and autoimmune disorders.

The prediction of protein complexes has advanced considerably with the advent of AlphaFold2(2), originally designed to predict the structures of individual proteins(3). However, AlphaFold2 accuracy decreases for interactions lacking co-evolutionary signals, particularly in complex biological systems such as host-pathogen(4), and for antibody-antigens(5). Recognising this limitation, two primary strategies have been established to improve prediction accuracy.

The first strategy involves extensive sampling methods demonstrated during the CASP15 competition(1; 6), where multiple structures were generated, increasing the likelihood of identifying accurate conformations. Several alternative methods have been proposed to increase sampling efficiency, including using dropout(1; 6; 7), masking columns in the MSA(8), and subsampling the MSA(9).

Several scoring functions have been developed to estimate the accuracy of predicted structures, starting with the introduction of pDockQ, used to predict the DockQ from AlphaFold2.0(3). With the introduction of AlphaFold-multimer(10), an internal score (ipTM) can be used to estimate the quality of the predicted interaction. Internally, AlphaFold2 and the other methods use the ranking confidence, a linear combination of 80% ipTM and 20% pTM. Several attempts have been made to improve the ranking. For instance, pDockQ2 (Zhu et al. 2023) can be used to predict the quality of interactions of each chain, actifpTM (Varga, Ovchinnikov, and Schueler-Furman 2024) is tuned to rank interactions involving flexible regions, and ipSAE (Dunbrack 2025) is developed to better separate true from false interactions than ipTM. All of these methods might provide an improvement over AlphaFold2 (i)pTM scores. However, it has not been clearly demonstrated that they significantly improve the per-target ranking of antibody-antigen complexes.

In this study, our objective was to evaluate new structure prediction methods for predicting antibody-antigen complexes using a new benchmark. We assess whether larger sampling, selection, and confidence metrics can improve the identification of accurately predicted structures.

## Material and Methods

### Dataset

The dataset consists of 110 antibody-antigen structures with less than 80% sequence identity to any protein in the training set. The dataset was created using a slightly modified version of the Antibody Antigen Dataset Maker (AADaM) (11), available from https://github.com/samuelfromm/AADaM-fork. We initially used all entries deposited in SAbDab (12) as of 2024-10-23. These structures are then divided into two subsets: a “before” dataset (2021-09-30) and an “after” (benchmark) dataset (2021-09-30). The chosen date, 2021-09-30, serves as the cutoff for the training data used by all benchmarked methods. The “before” dataset was filtered to retain only structures in which at least one antigen chain is a peptide or protein.

Antibody-peptide structures were excluded from the benchmark dataset, which was further filtered to include only structures determined by X-ray diffraction or electron microscopy, with a resolution of 3.5 Å or better. Structures with nonstandard residues were excluded unless at the sequence ends, in which case they were trimmed. Structures missing more than 10% of residues were also discarded. Structures are discarded from the benchmark dataset if the sequence identity between the benchmark and “before” dataset exceeded 80%. Sequence identity between the two datasets was calculated individually for the heavy (H), light (L), and antigen chains using a global alignment. The benchmark set itself is further clustered by sequence identity using the same procedure and threshold. During clustering, the structure with the fewest missing residues in the H, L, and antigen chains is retained. If two structures have the same number of missing residues, the one with the shorter antigen sequence is selected. One of the now 111 complexes failed due to an HHblits-related issue and was therefore excluded, leaving the final dataset with 110 proteins. In the final set, no two Antibodies are identical, while one antigen is present twice, bound to two different antibodies.

We also quantified how the benchmark size would change if redundancy was assessed using concatenated complementarity-determining regions (CDRs) sequences. Applying the same 80% identity threshold retained 416 complexes, reflecting the much higher diversity of CDR regions compared to full antibody sequences. Furthermore, we examined CDR redundancy in 110 benchmark complexes relative to the training set and found that although some loops are identical to those in the training set, no single antibody has all its loops identical to any antibody in the training set. The maximum sequence identity for the concatenated loops is 72%. For antigens, 15 sequences were found in the training set, while the other 110 are unique to the benchmark (note that some complexes have more than one antigen chain). In Figure S1, it can be seen that most antibodies share a maximum identity to a member of the training set of about 50% for the concatenated loops, although individual loops have higher identities.

### Structure Generation

#### AlphaFold2.3 methods to enhance sampling

To increase the sampling diversity, we examined previously described methods (6; 8; 9) by modifying the AlphaFold2.3.2 code. All methods were run without relaxation. The maximum template date is set to 2021-09-30. We used the latest version of the weights (v3) and for each of the five available network models, 40 predictions were generated, resulting in a total of 200 models per method.

In total, five different AlphaFold2-based methods were evaluated:

- **AlphaFold2.3**: AF2.3 with default parameters.
- **AF2.3-SubSampl-32-64**: AF2.3 with MSA subsampling using `num_msa=32` and `num_extra_msa=64` compared to the default values of 508 and 2048, respectively.
- **AF2.3-SubSampl-16-32**: AF2.3 with MSA subsampling using `num_msa=16` and `num extra msa=32`.
- **AF2.3-Dropout**: Using the ideas in AFsample (6), i.e. we turn on dropout during interference.
- **AF2.3-ColMask**: Using the ideas in AFsample2 (8), i.e. we randomly mask 15% of the columns in the MSA.

The code for our implementation of the above methods is available at https://github.com/samuelfromm/alphafold-clone. It should be noted that we used the default parameters for all methods. Other hyperparameters, such as dropout rates, might yield better performance. In CASP, up to five models were allowed, and it is often good to submit a diverse set of models (Elofsson 2025), so if such a strategy were applied here, the difference to the best possible model would decrease. However, the best approach for selecting five models has not been well studied; therefore, we have focused only on selecting the top-ranked model.

#### AlphaFold 3, Chai-1 and Boltz-1 models

For each complex in the dataset, 40 random seeds (1-40) were used to generate five models each, resulting in a total of 200 models per complex.

#### MSA generation

Multiple sequence alignments (MSAs) were generated using the AlphaFold 2.3.2 pipeline (specifically commit `f251de6`). For running AF3 and Boltz-1, we specifically used the `bfd_uniref_hits.a3m` file produced by this pipeline, while for Chai-1, MSAs were generated using the `--use-msa-server` option via the ColabFold (13) server.

### Evaluation

#### Scoring

To calculate scores, we created a Snakemake workflow. For each prediction, the corresponding predicted model (query) and the PDB ground truth structure of the corresponding PDBID (reference) were used as inputs.

AADaM(11), the pipeline used to generate our dataset, outputs a FASTA sequence for each antibody-antigen structure. However, these structures do not always align fully with their corresponding PDB entries. Since AADaM focuses on the antibody-antigen interface, some chains present in the original PDB file may be excluded from the interface structures used in our dataset. Additionally, the sequence in the reference structure from the PDB is often shorter than the SEQRES sequence used to generate the protein structures. To address both issues, we first used MMalign (14) to calculate the alignment between the query and reference structures. We then take each pair of aligned chains and calculate a sequence alignment between them. Based on this sequence alignment, we select the residues present in both the query and the reference structure.

Using the cropped query and reference structures as input, we calculate several metrics:

1. We used MMalign to calculate the TM-score (15).
2. We calculate the DockQ score (16; 17) for the antibody-antigen interface (“merging” multi-chain antigens into a “single chain”). This results in a single DockQ score for each complex.
3. Similarly, we calculate the ipSAE score (18) with a PAE and distance cutoff of 10 Å for the antibody-antigen interface, resulting in a single score for each complex.
4. We calculate pDockQ version 2(19).
5. We extract the confidence scores from the predictions as output by the respective method: `ranking_confidence` for AlphaFold-based methods, `confidence_score` for Boltz-1, and `aggregate_score` for Chai-1.
6. We calculate the aligned error between the cropped query and reference structure as well as the aligned error ranking confidence (see section *Aligned Error* ).

The workflow is the same for all samples and methods, except for the last step: the aligned error analysis was performed only for a selection of methods. The complete workflow is included in the provided code.

In some cases, MMalign cannot properly align the chains, indicating that the query and reference structures are dissimilar and that the predicted structure is of very low quality. We still run the rest of the workflow even when the number of aligned residues is extremely low. When the number of aligned residues, normalised by the query length, is below 0.9, the resulting DockQ scores typically fall between 0 and 0.05. We consider these low scores to accurately reflect the poor quality of the predicted interfaces in such cases.

### Sampling

For each method, we generated 200 models for each complex and treated them as independent predicted structures, although in AF3’s inference, each random seed generates 5 different models. These models use the same information as the pairformer, but differ in the built-in randomness of the diffusion process for training. To assess the impact of sampling, we randomly select k models with replacement, identify the best (highest DockQ) and top-ranked (highest confidence) models, and record their DockQ scores. For each sampling size k, we average the recorded DockQ score over 50 iterations. We then average the resulting score over all IDs and sample sizes.

### Structural Comparison

We used both global and interface similarity to investigate whether structural similarity could be detected between the proteins in the test and training sets. First, we used FoldSeek-Multimer (20; 21) to identify global similarity between a complex in the test set and any protein present in PDB prior to the split date (2021-09-30). For each complex, the maximum TM-score(15) was normalised by the length of the query protein. Next, we created a smaller dataset to examine the interface similarity using USalign (22). This dataset included all 1998 PDB entries with a TM-score > 0.5 to any complex in the test set. We first extracted all interface residues from the proteins in the test set to evaluate interface similarity. Interface residues were defined as any antigen residue containing an atom within 12 Å of the antibody or vice versa. Next, the interface was aligned to the 1998 PDB entries mentioned above using USalign with the `-mm 1` flag, and the maximum TM-score was recorded.

### Model diversity, PconsDock

To estimate the diversity of models for a given target, we used PconsDock(23), i.e. a pairwise comparison of all the models for a target using DockQ. Due to computational constraints, we did not pairwise compare all 200 models. Instead, for each model, we randomly selected 20 other models for comparison. We then report the average DockQ per target or per model.

### Aligned Error

AlphaFold is trained to estimate two types of errors: the pLDDT, which estimates the error for each residue, and the PAE, which estimates the *Aligned Error* for each pair of residues. The predicted Aligned Error (PAE) is then used to calculate pTM and ipTM, and hence the confidence score. We want to examine if it is the quality of these estimates or the functional form of ipTM/pTM itself that hampers discrimination among model predictions. To test this, we evaluate discrimination performance with oracle-based AEs (aeTMs) for a given prediction, described below.

We follow the notation in (2) [§1.9.7]. Assume that we are given two structures - a reference and query structure - for the same sequence. Let 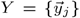 and 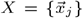 denote the *C*^*α*^-positions of the reference and query structure, respectively, let *T*_*Y,i*_ and *T*_*X,i*_ be the corresponding backbone frames, and let 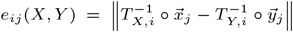 be the error in the *C*^*α*^-position of residue *j* when the query and reference structures are aligned using the backbone frame of residue *i*. The non-symmetric matrix 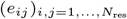, where *N*_res_ is the number of residues, is called the aligned error (AE) between the two structures. Let

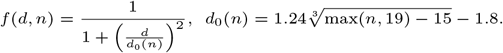

We define the aligned error TM-score (aeTM) and aligned error interface TM-score (aeiTM) as

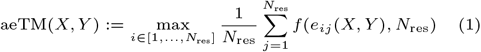

and

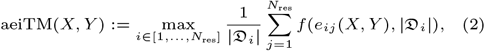

respectively. Here, *𝔇*_*i*_ denotes the set of all residues except residue *i*

Note that the aeTM score and the aeiTM scores can be viewed as idealised versions of the predicted TM (pTM) score and interface predicted TM (ipTM) score, respectively. Indeed, let us replace *X* with a predicted structure and assume that the ground truth structure *X*^*true*^ is unknown. Assuming that we have a random variable *Y (ζ)*, sampling structures from a distribution of probable ground truth structures. Then the pTM and ipTM scores are given by

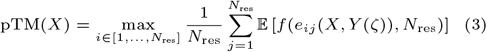

and

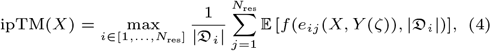

respectively (see AF2, §1.9.7 and AF2-multimer, §7.9). Here, the expectation is taken over the probability distribution of *e*_*ij*_ *(X, Y* (ζ)). Now, in an ideal situation, ℙ (*Y* (ζ) = *X*^*true*^) = 1, i.e. the random variable *Y* takes on the values *X*^*true*^ with probability 1, in which case 𝔼 [*f* (*e*_*ij*_ (*X, Y* (ζ)))] = *f* (*e*_*ij*_ (*X, X*^*true*^)) so that pTM(*X*) = aeTM(*X, X*^*true*^) and similarly ipTM(*X*) = aeiTM(*X, X*^*true*^).

In practice, AlphaFold learns to predict the probability distribution of the elements of the aligned error *e*_*ij*_ (*X, Y* (ζ)). To this end, the distribution of *e*_*ij*_ is discretised into 64 bins, ranging from 0 to 31.5 Å with a bin width of 0.5 Å, where the final bin captures any errors larger than 31.5 Å. When calculating the aligned error between two structures *X* and *Y*, we do, however, allow unlimited error values.

While the pTM score is a predicted metric, the aeTM takes as input any two structures of the same sequence, similarly to the TM-score(15). Note also that the aeTM score is bounded from above by the TM-score (see AF2, §1.9.7).

Finally, recall that AlphaFold’s ranking confidence score is defined as 0.2 *·* pTM +0.8 *·* ipTM. In a similar fashion, we define the aligned error ae ranking confidence (aeRankConf) as 0.2 *·* aeTM +0.8 *·* aeiTM.

## Results and Discussion

Here, we evaluated the performance of AlphaFold2.3 (10), AlphaFold3 (24), Boltz-1 (25), and Chai-1(26), using a dataset of 110 unseen antibody-antigen structures with limited identity to the training set, for varying numbers of generated protein models. For AlphaFold2.3, we also examined other strategies reported to increase sampling efficiency.

Several studies have reported that generating more samples improves the quality of AlphaFold2 predictions (6; 7; 27). Further, in the AlphaFold3 (24) paper, it is claimed that generating thousands of samples improved the predictions of antibody-antigen complexes. We generated 200 models for each method and examined their performance, measured by the DockQ score(16; 17) between the antigen and the antibody. The performance was measured for each sampling size by using the mean DockQ of the best model for each target and the mean DockQ of the top-ranked models, see Methods for details. A summary of the results is presented in Figure 1.

**Fig. 1.**
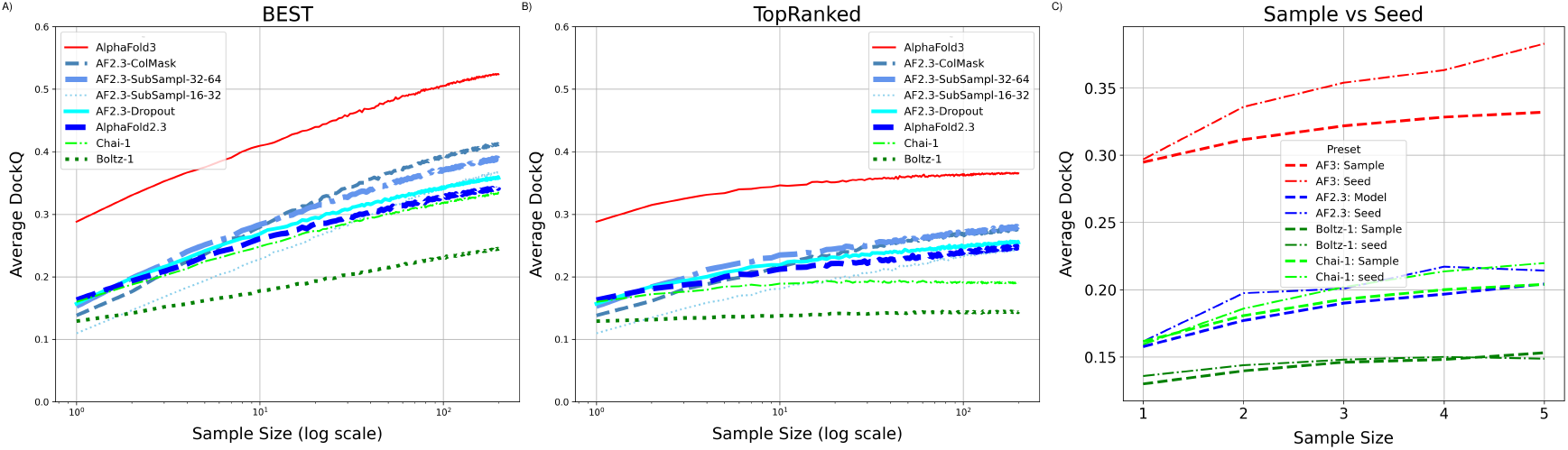
For each method and sample size *n* we randomly select *n* proteins out of the 200 structures we created (with replacement). Out of these *n* selected models, we select the model with the highest DockQ score (A) or highest ranking confidence (B), respectively, and record its DockQ score on the y-axis. C) Identification of the best, i.e. highest DockQ score, model when changing either the seed or only the sample for up to five models

### AlphaFold3 makes better predictions than the other methods

First, we will discuss the ability to generate a good model, assuming an oracle exists to identify the best model among all models generated by a method. Figure 1A shows that AlphaFold3 outperforms all other methods by some margin, while AlphaFold2, Chai-1, and Boltz-1 show similar performance when a single model is generated. One example where AF3 performed considerably better than other methods is for the target 8TQ7, see Figure 5. Here, the AF3 prediction is spot-on, while all other methods predicted binding to different regions of the antigen.

Secondly, it can be observed that all methods produce better protein models as sampling increases. A roughly linear increase is observed with the logarithm of the sample size. This means that the average DockQ for AF3 of the best model increases from below 0.3 to above 0.5 when 200 samples are generated. A similar increase is observed for AlphaFold2 and Chai-1(26), while a smaller improvement is observed for Boltz-1.

In AlphaFold3, additional models can be generated by rerunning the entire pipeline with a different random seed or by letting the diffusion model produce multiple models. Figure 1C shows that using different seeds yields greater structural diversity than generating more samples from the diffusion step alone.

### AF2.3-ColMask (AF-Sample2) is the most efficient strategy to increase sampling

As shown before, it is possible to increase the diversity of the generated proteins using several different strategies. We examined these strategies by modifying AlphaFold2.3, as the AlphaFold3 license does not allow such modifications and because this study was initiated before the release of AlphaFold3. In CASP15, it was shown that increased sampling via dropouts improved prediction accuracy, particularly for antigen-antibody pairs (6) using AFsample. We refer to it here as AF2.3-Dropout, as the implementation is not identical. More recently, the Wallner group developed AFsample2 (hereafter referred to as AF2.3-ColMask), an even more aggressive method to add noise by masking entire columns in the MSA (8). It is also possible to generate alternative conformations by subsampling the MSA(9).

Figure 1A shows that, on average, all methods generate slightly worse models than the default AlphaFold2.3 setting when only one model is generated, see `SampleSize` = 1(10^0^). The difference is negligible for all methods except for AF2.3-ColMask (which is very similar to AFsample2), which produces significantly worse average models, DockQ of 0.138 vs 0.163. However, as more models are generated, all these methods produce better models than AlphaFold2.3 does natively. Overall, AF2.3-ColMask generated the best protein models, likely due to its more aggressive noise addition.

### A large gap between the best and top-ranked models

It is necessary to identify good models as well as generate them. Figure 1B shows the average DockQ score for the top-ranked model against sample size. For all methods, the increase in average DockQ with sample size is much smaller than what is observed for the best model (Figure 1A). This means that the internal ranking of the models cannot identify the best generated model. For AlphaFold3, it means the average DockQ increases only from 0.29 to 0.37, rather than 0.52. For Boltz-1 and Chai-1, there is hardly any improvement with larger sample sizes (from 0.12 to 0.14). For the AlphaFold2.3-based models, a similar trend to that observed for AlphaFold3 is evident. Here, AF2.3-SubSample performs as well as AF2.3-ColMask when 30 or more models are generated. It is clear that better detection of top-ranked models would significantly improve the prediction of antibody-antigen structures.

In Figure 2, we take a more detailed look at the distribution of the scores for each target. It can be seen that for all methods, the targets can, in general, be divided into three types. To the left in the plots are the easy-to-predict targets, where most models are correct, though a few are incorrect. To the right in the plots are challenging targets, where most models are wrong, but a few good models exist for some targets. In the middle are models with many good and bad models. Further, the main difference between the four methods is the number of targets within each category, i.e. AlphaFold3 has many more easy-to-predict targets.

**Fig. 2.**
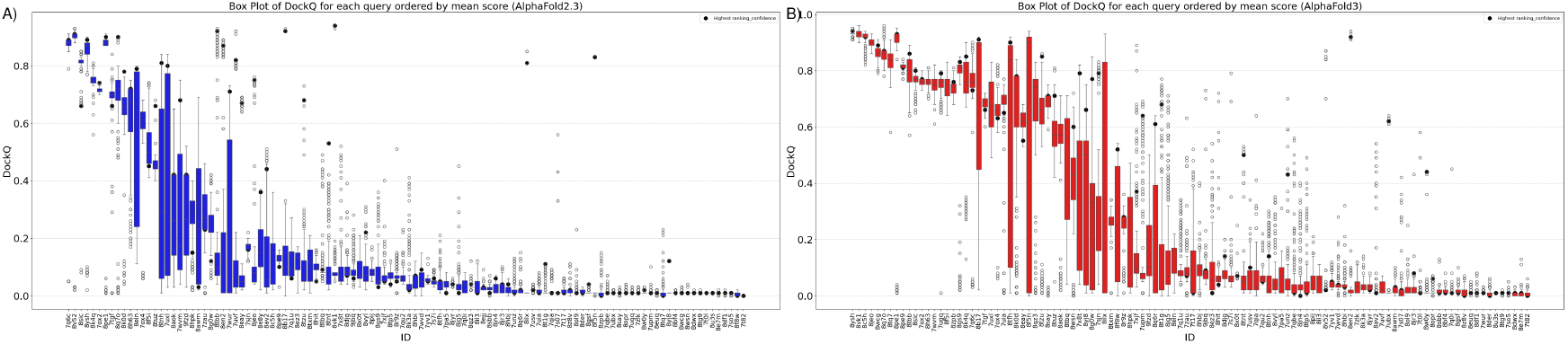
Each figure shows a box plot for the DockQ score for each target ID, sorted by mean DockQ score for that target ID and that method. The highest-ranked model is highlighted with a thick dot. A) AlphaFold2.3, B) AlphaFold3, for Boltz-1 and Chai-1 see Figure S2. It can be seen that the variance of DockQ and ranking confidence is highest for targets with intermediate difficulty (see Figure S3 and S4, and that these two variations correlate for AlphaFold3, Figure S5 and S6

In Figure 3, it can be seen that for both top-ranked and best models, some targets where AlphaFold2.3 outperforms AlphaFold3 exist. This indicates that there is no unique set of easier models; instead, there is a larger set of models where AlphaFold3 produces significantly better structures than the other methods.

**Fig. 3.**
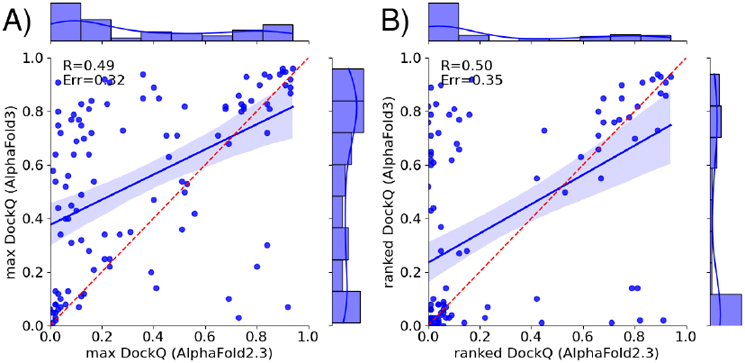
DockQ scores for AlphaFold3 vs AlphaFold2.3 for best models (A) and top-ranked models (B)

### Has AlphaFold3 learned to predict structure beyond its training set?

Next, we investigated how AlphaFold3 can predict many targets almost perfectly, while other methods failed for the same targets. There should be no coevolutionary signal between the antibody and the antigen; therefore, AlphaFold3 may have learned something about the physics governing the interactions. Alternatively, it may have identified patterns in the database that other methods did not. To answer this, we compared each target in our test set with the training data, i.e., all of PDB until Sep 2021. The search was made either by examining just the interface (Figure 4A) or the entire complex (Figure 4B). For each complex in our test set, we noted the maximum TM similarity to any complex in the training set for either the entire complex or just the interface region.

**Fig. 4.**
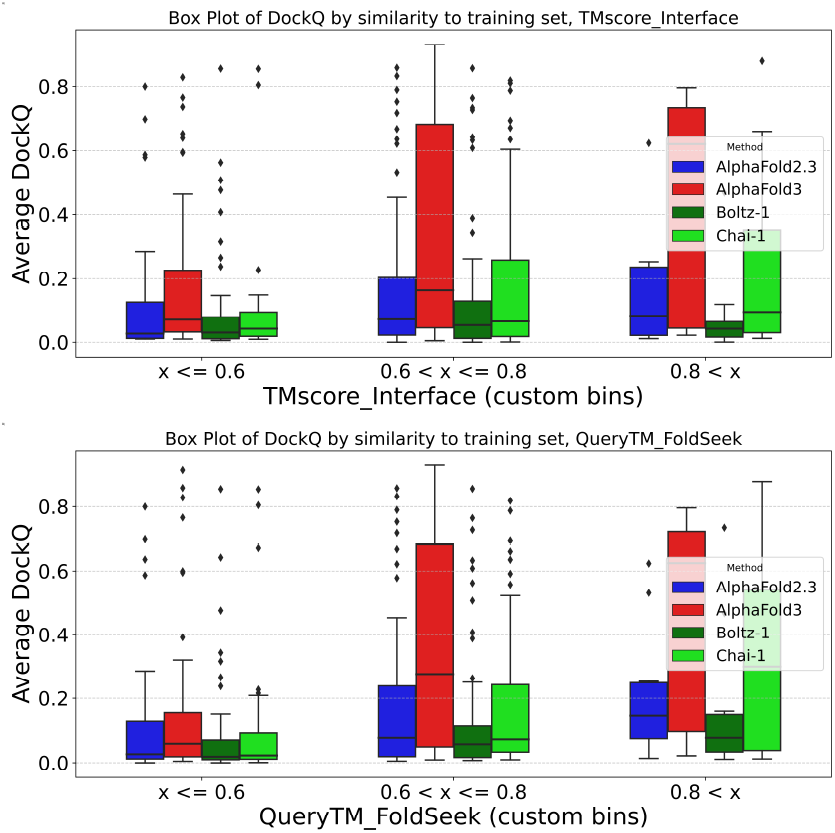
Prediction performance vs. similarity to the training data for AlphaFold2.3 and AlphaFold2 using Interface TM-score similarity (A) or Query TMscore similarity (B) vs the mean AbAG-score for each target. For Max or First ranked scores, see Figure S7

As shown in Figure 4, the mean DockQ score significantly increases when a structurally similar antibody-antigen complex exists in the training set of AlphaFold3, and to a lesser extent for Chai-1. When the best model for each target is used, a similar trend can be seen for all methods, Figure S7. However, it should be noted that well-predicted models exist in which no structural similarity can be detected, and that the prediction is often dire, even if a structurally similar complex is present in the training set, as seen in Figure S8. In general, the correlation between the AbAG-DockQ score and the similarity to the training set is low (Cc=0.32 for complex search and 0.17 for interface) (Figure S9), further supporting the idea that similarity with the training set is neither a requirement nor a guarantee for a correct prediction. We could not detect any trends for better performance for targets with high sequence identity in either the CDR regions or for the antigens, Figure S10 and S11.

We also noted that it is not trivial to understand what AlphaFold3 might have used from the training set to make excellent predictions. Figure 5B shows an example of the most similar hit in the training set to a model that AlphaFold3 (but not other methods) predicted well (8TQ7). It can be seen (in magenta) that the best hit (6DDM) has a similar beta-sheet domain that binds to the antibody. However, it is also clear that the interface is not identical (the interface TM-score(15) is 0.63), indicating that, in this case, AlphaFold3 may have learned the orientation of the domains rather than the exact packing of the interface.

**Fig. 5.**
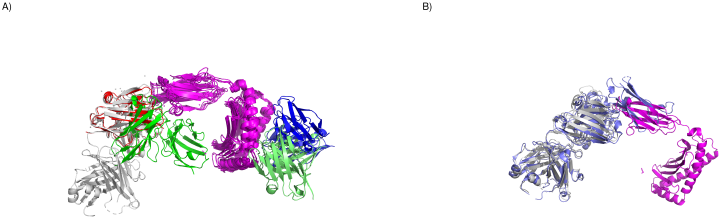
A) Example (8TQ7) of a prediction where AlphaFold3 is superior to all other methods. The antigen is shown in Magenta, Grey - PDB structure, Red AF3, Green Boltz-1, Lime Chai-1, Blue AF2.3. B) Superposition of 8TQ7, in magenta and grey, with the most similar protein complex in the training set (6DDM) in blue.

### Ranking of AlphaFold3 models

Figure 1 clearly shows that an improved ranking of the models for each target would significantly improve antibody-antigen predictions. The average DockQ score would increase from 0.37 to 0.52. While the correlation between ranking confidence and DockQ ) across all targets is reasonably good (Cc=0.80 for ranking confidence), it is quite poor when considered on a per-target basis (mean CC=0.28), as can be seen in Figure 6. Although for some targets ranking confidence and DockQ correlate, for many they do not. Some targets have almost identical ranking confidences, but the DockQ varies widely, while others, including 7WVM, have virtually identical (bad) DockQ scores, while the ranking confidence differs widely. Alternative measures, such as pDockQ, which are also based on predicted qualities (pLDDT and PAEs), show very similar behaviour (data not shown).

**Fig. 6.**
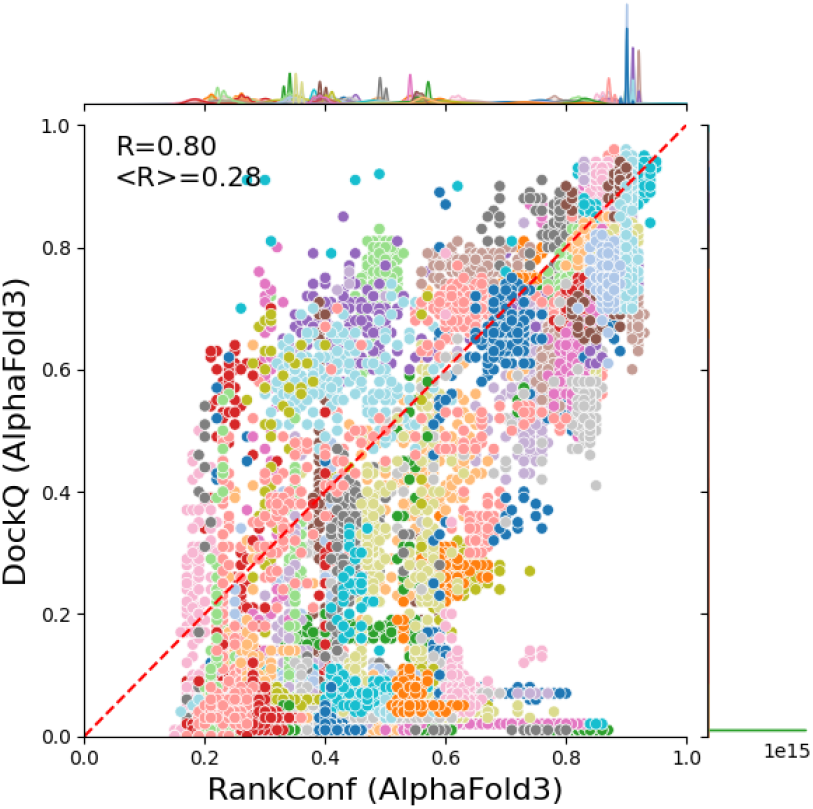
DockQ scores vs Ranking Confidence for all AlphaFold3 models. All models are shown in a single colour for each target. The overall correlation coefficient is 0.8, while the mean correlation coefficient per target (*< R >*) is only 0.28. . A plot with a maximum of ten targets per plot is shown in Figure S15

An alternative method to estimate the quality of a model is to use what we earlier referred to as PconsDock(23), i.e. to pairwise compare all the models for a target using DockQ. In Figure S12 and S13 we show that the PconsDock score correlates as well as ipTM with the real DockQ scores, but is no better at ranking the targets. The reason is, obviously, because for a hard target, the models do not agree on how to locate the antibody on the antigen. In contrast, for an easy target, all antibodies are placed at the same location, as shown in Figure S14. Interestingly, there is one target, 8U3S, where both ipTM and PconsDock scores are high, although the DockQ score is low; see Figure S14. In this case, SabDab is likely wrong, as the asymmetric unit does not correspond to the biological assembly according to PDB.

### Ranking analysis

One of AlphaFold2’s most innovative features was its ability to accurately estimate the quality of the generated protein structures. The estimates stem from two predicted features: the prediction of the accuracy of an individual residue (pLDDT) and the Predicted Aligned Error (PAE), which estimates the error in the distance between pairs of residues (2). Since the introduction of AlphaFold-multimer, these estimated distances have been used to calculate ipTM, specifically estimating the accuracy of interactions between chains.

Attempts to improve these scores have been proposed since the introduction of pDockQ (3), which predicts the DockQ score for protein models generated by AlphaFold2.0. pDockQ bases its predictions on the number of interacting residues and their pLDDT values. Several other methods based on similar ideas have been introduced later. Recently, attempts have been made to improve AlphaFold’s ipTM and pTM scores (and thus the ranking confidence) by recalculating them from PAE values (18; 28). These methods modify the functions (3) and (4) used to calculate the pTM and ipTM scores, respectively, while still using the same features used to calculate the ipTM and pTM scores.

To demonstrate the importance of accurate PAE predictions, we have also included these measures calculated from perfect PAE values, aeTM and aeiTM. One example of how this affects predictions is shown in Figure 7A-D. The two models are of 8HIT, one is almost perfect (DockQ: 0.690), while the other is rather poor (DockQ: 0.040). However, the PAE matrices for the two cases are almost identical.

**Fig. 7.**
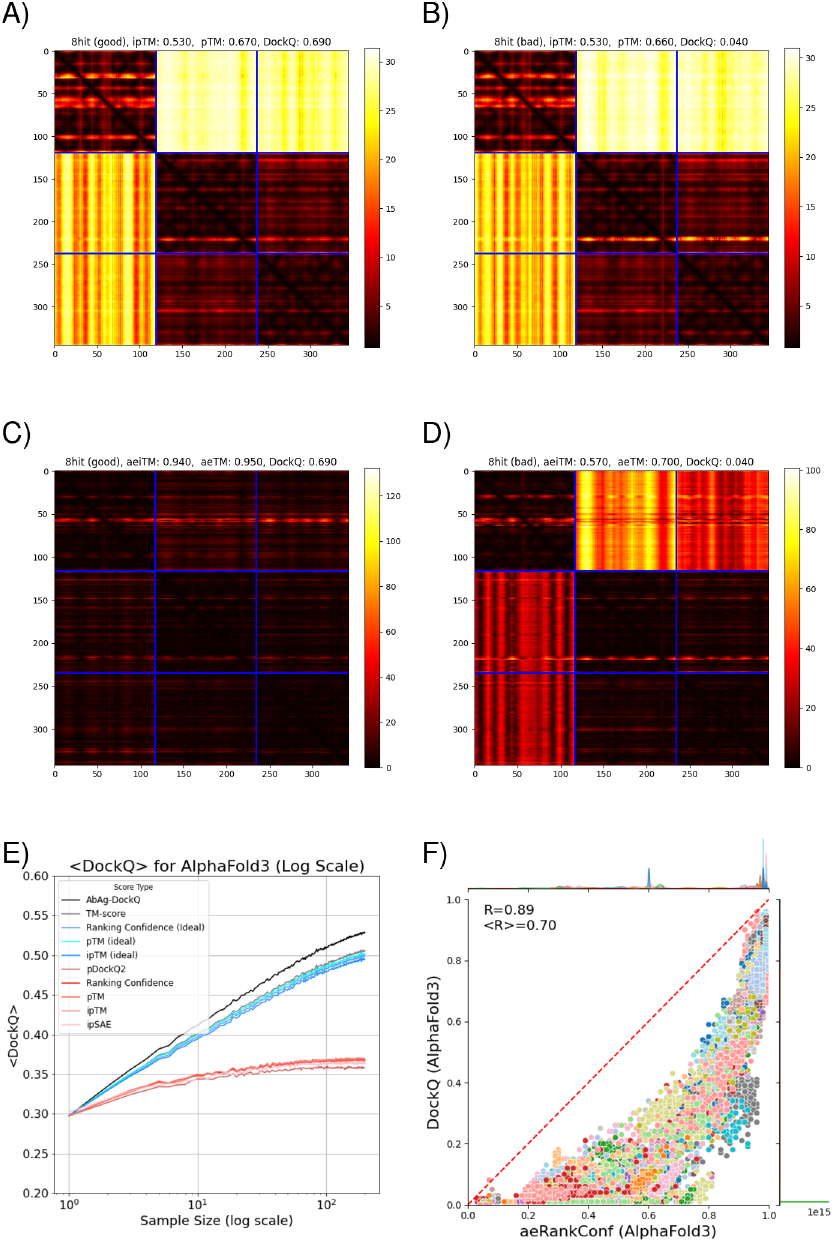
A) and B) PAE plots for two models of 8HIT. The chain breakers are marked in red. The mean absolute error (mae) between the two PAE matrices is 1.03 across the whole matrix and 1.37 across the inter-chain region. C) and D) AE plots for the same two models 8HIT. The mean average error between the two AE matrices is 13.24 across the whole matrix and 19.53 across the inter-chain region. E) Ranking using different ways to use the PAEs to calculate the scores, in red methods using predicted PAEs and in blue using AE (i.e. idealised PAE) scores. The primary issue is that the ground-truth aligned error values are not sufficiently accurate, resulting in a significant performance gap. F) the aligned error ranking confidence values (compare with Figure 6).

To better understand the impact of feature quality versus function accuracy, we investigate how well the ideal aligned-error scores, aeiTM and aeTM (see equations (1) and (2)), perform. We compared these measures on their ability to identify top-ranked model, as well as their correlation with DockQ, both overall (R) and per target (< R >), see Figure 7E and Table1. For measures based on predicted PAE values - regardless of the specific choice of measure - (i) the average DockQ score of the top-ranked model is significantly lower than the best possible (DockQ=0.35, vs 0.54), (ii) the overall correlation with DockQ is acceptable (R > 0.7), while (iii) the per-target correlation is much worse (< R >< 0.30). At the same time, using any of the measures with idealised PAE predictions solves these problems. Figure 7E shows the average DockQ score for each ranking method and sample size. Selecting by aeiTM yields results much closer to the best possible outcome compared to selection by ipTM, indicating that the limiting factor when it comes to the ipTM score, at least for antibody-antigen complexes, is the accuracy of the predicted aligned error (and not necessarily a problem with the way the ipTM score is calculated itself). In Figure 7F, it can be seen that the average per target correlation is almost as high as the overall correlation (R=0.70 vs 0.89).

**Table 1.** Summary for different methods to rank AlphaFold3 models, Average DockQ (< DockQ >) and Spearman correlation for all targets (R) or average per target < R >.

| Method | $< DockQ >$ | R | $< R >$ |
| --- | --- | --- | --- |
| DockQ | 0.544 | 1.000 | 1.000 |
| aeTM | 0.516 | 0.814 | 0.665 |
| aeiTM | 0.508 | 0.873 | 0.562 |
| aeRankConf | 0.511 | 0.879 | 0.595 |
| pTM | 0.369 | 0.670 | 0.200 |
| ipTM | 0.370 | 0.707 | 0.220 |
| RankConf | 0.375 | 0.799 | 0.214 |
| pDockQ2 | 0.352 | 0.603 | 0.184 |
| ipSAE | 0.353 | 0.750 | 0.247 |

### Analysis of the PAE matrices

As can be seen in Figure 7 the PAE matrices often have some sort of patterns. To examine whether this could have a rational explanation, we compared all PAE matrices across models. First, we defined two regions of the PAE matrix. Region1 is the region to the top right related to the antibody-antigen contacts, while region2 is the corresponding area in the bottom left part of the matrix. Region1 is related to the PAEs when the models are aligned on the antigen, while Region 2 is related to the PAEs when the alignment occurs on the antibody.

First it can be noted that the scores are almost always lower (better PAEs) in Region2 than in Region1, this is likely just an effect of size. However, this results in the fact that in almost all cases pTM and ipTM are calculated from one line in this region. Secondly, the correlation between different targets within the same region is much higher than that between region1 and region2 (Figure S16), indicating that it might be of important to develop a score that takes both regions into account.

Next we analysed the correlation between rows and columns in the two regions, see Figure S17, this refers to how consistent the PAEs are when aligning on a specific residue in the other chain. It can be seen that the strongest correlation is for rows in region2. This means that the strongest pattern occurs when the antigen is aligned with antibody residues, i.e. it is not particular residues (loops) in the antibody that dominate the patterns. We observed that the lowest PAEs were often found on surface exposed residues (data not shown).

## Conclusion

In this study, we benchmarked different sampling and ranking strategies against a selected ensemble of unseen antibody-antigen complexes.

Our findings reveal that (i) AlphaFold3, but not Chai-1 and Boltz-1, consistently offers improved predictions over AlphaFold2, (ii) larger sampling sizes correlate with an increased probability of generating at least one good model, (iii) a crucial bottleneck persists in accurately identifying the best model among the generated models, and (iv) AlphaFold3’s performance drops considerably for complexes without prior structural similarities to the training dataset, underscoring the model’s dependence on previously encountered data for accurate predictions, the same is not seen for analysing sequence similarity to the training set. However, even if we see an increased success rate for targets with greater structural similarity to the training set, it is clear that this is neither a sufficient nor a necessary condition, as there are successful predictions with low similarity and unsuccessful predictions with high similarity.

Furthermore, to analyse the confidence metric used by AlphaFold (ipTM score), we introduced an idealised version of the ipTM score, showing that if the underlying estimates of accuracy (the predicted aligned error) are sufficiently accurate, the default ipTM formula is sufficient to achieve close to ideal per-target ranking and thereby significantly increase the DockQ score of the top-ranked model. This indicates that the limitation lies not in the ipTM formula itself but in AlphaFold’s ability to accurately predict aligned errors.

## Author contributions

S.F. prepared the dataset. S.F. and M.L. predicted all the structures. A.E. and S.F. performed the analysis. A.E. wrote the initial draft of the manuscript. All authors contributed to the final version of the manuscript.

## Achnowledgement

AE was funded by VetenskapsrÅdet, Grant No. 2021-03979, and the Knut and Alice Wallenberg Foundation, Grant No. 2022.0032. The computations and data handling were enabled by the supercomputing resource Berzelius, provided by the National Supercomputer Centre at Linköping University, the Knut and Alice Wallenberg Foundation, and SNIC, grant numbers SNIC 2021/5-297 and Berzelius-2021-29.

## Supplementary Material

**Fig. S1.**
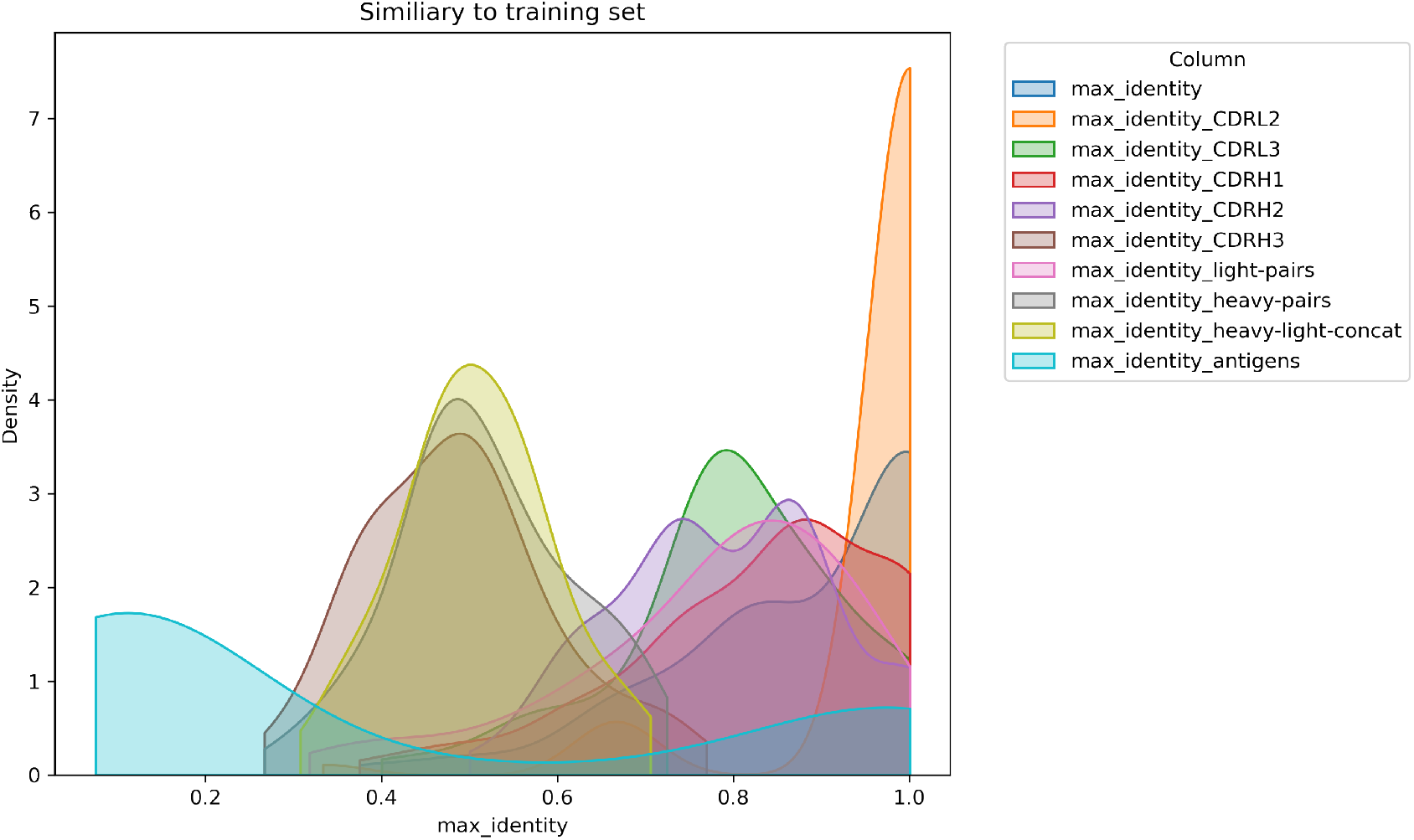
Distribution of maximum sequence identity to training set for individual loops in the antibodies, concatenation of all loops in heavy/light chains or both, as well as the antigen. For all individual loops, there are almost identical sequences in the training set. In contrast, for the concatenated loops, the identity is about 50%, and for the antigens, there is a bimodal distribution with a fraction of almost identical antigens in the dataset. At the same time, a majority has less than 30% sequence identity.

**Fig. S2.**
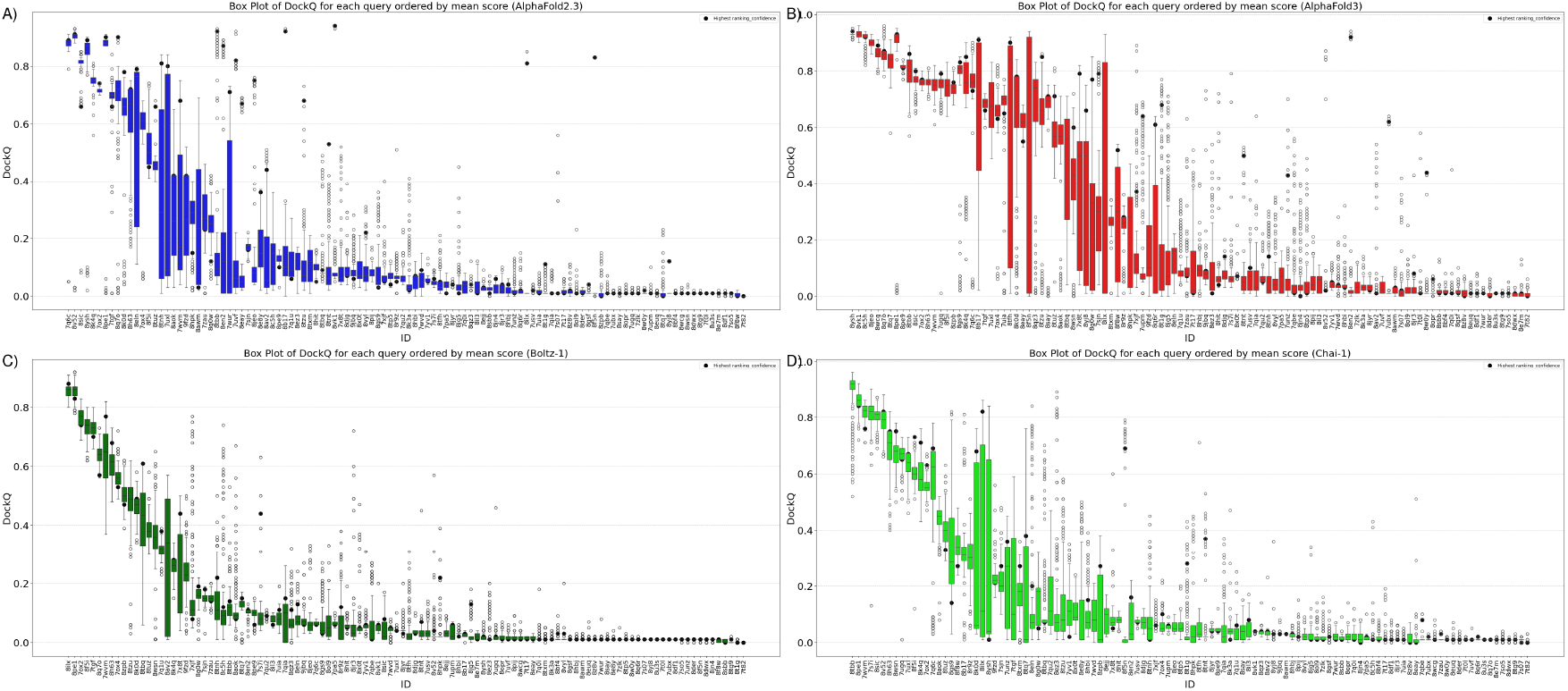
Each figure shows a box plot for the DockQ score for each target ID, sorted by mean DockQ score for that target ID and that method. The highest-ranked model is highlighted with a thick dot. A) AlphaFold2.3, B) AlphaFold3, C) Boltz-1, and D) Chai-1.

**Fig. S3.**
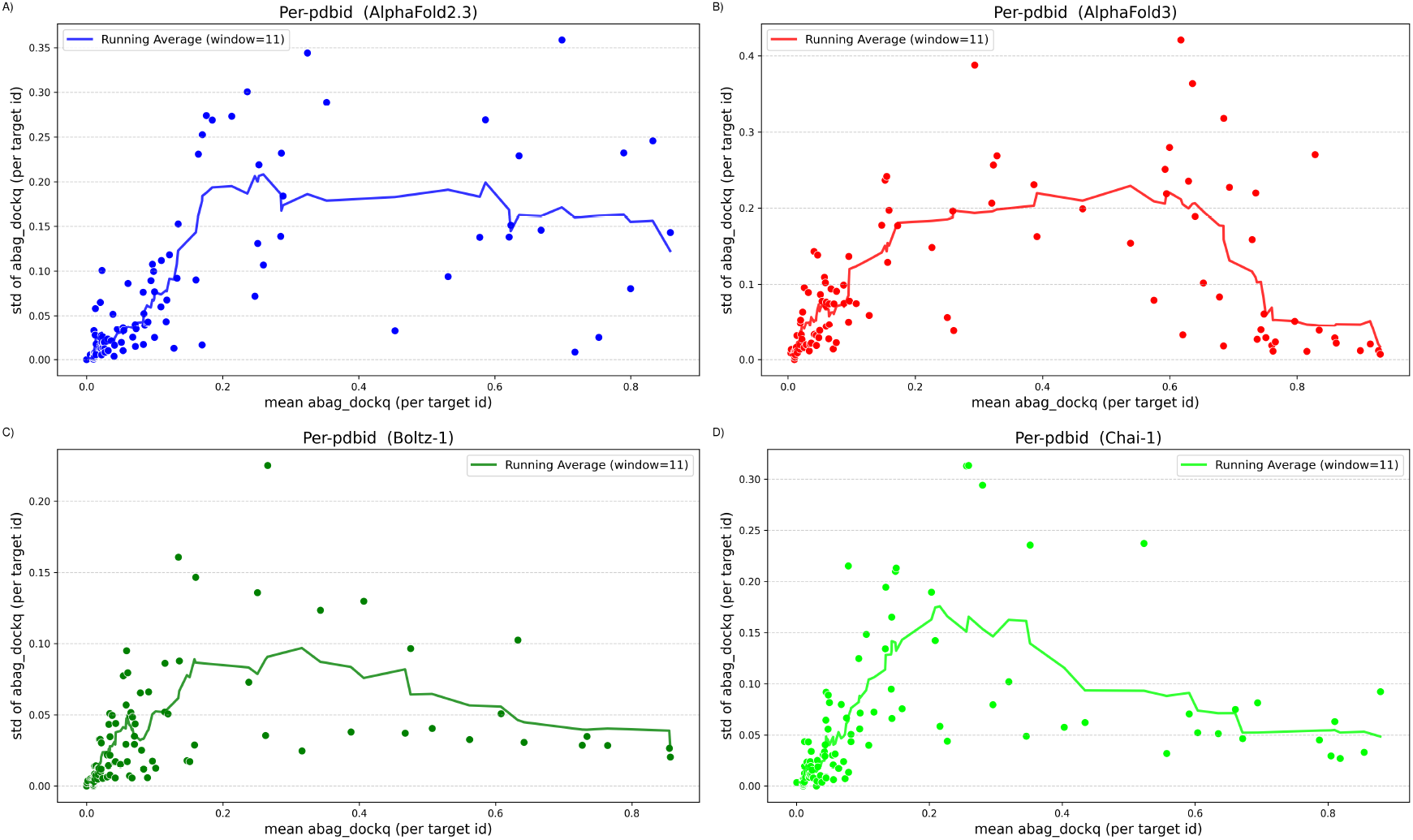
Each figure shows a scatter plot for the mean vs Standard Deviation of the DockQ scores for each target. A) AlphaFold2.3, B) AlphaFold3, C) Boltz-1, and D) Chai-1. The running average (over 11 sample) show a trend that the variation is largest for “intermediately” easy targets.

**Fig. S4.**
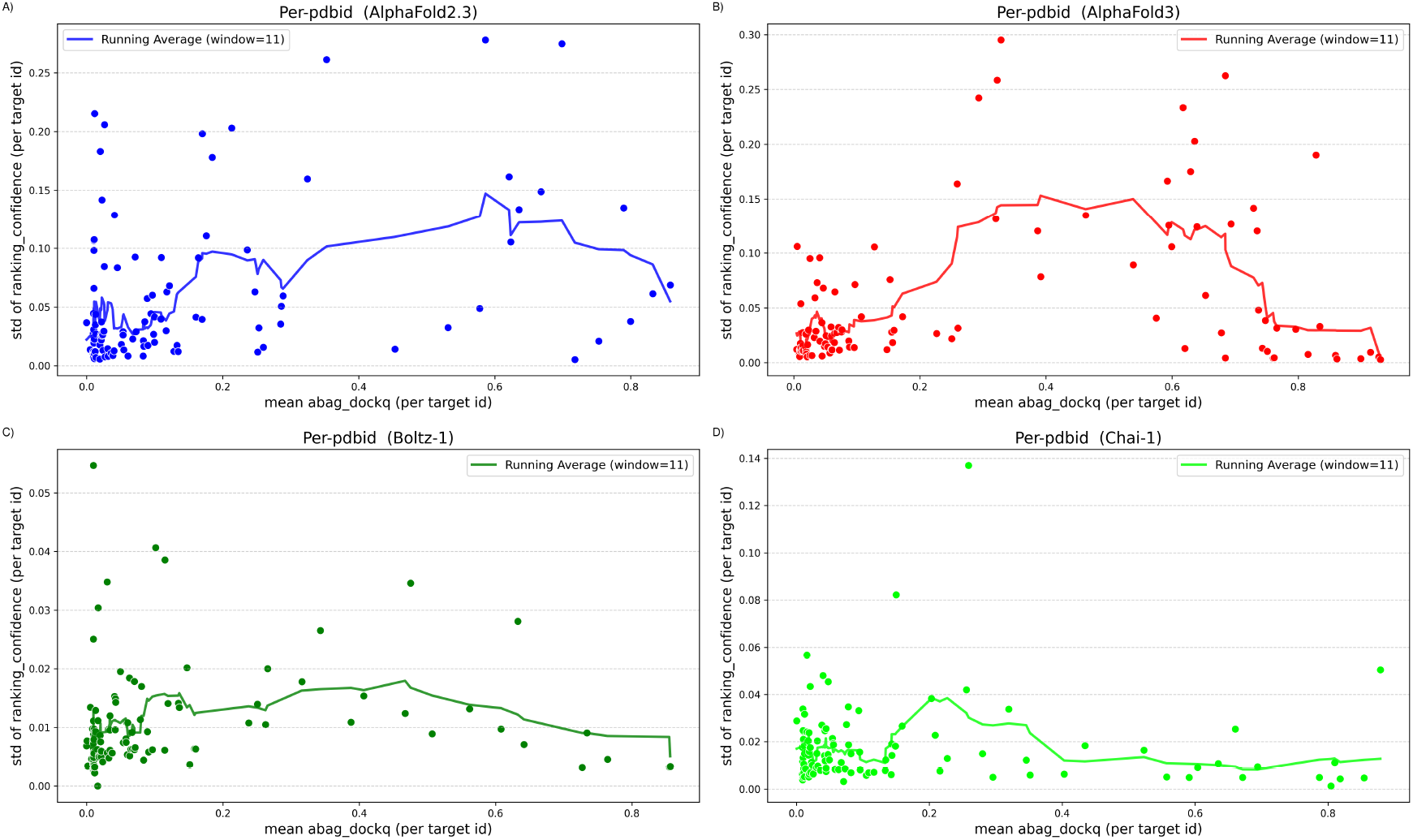
Each figure shows a scatter plot for Standard Deviation of the ranking confidence scores vs the mean DockqQfor each target. A) AlphaFold2.3, B) AlphaFold3, C) Boltz-1, and D) Chai-1. The running average (over 11 sample) shows a minor trend that the variation is largest for “intermediately” easy targets.

**Fig. S5.**
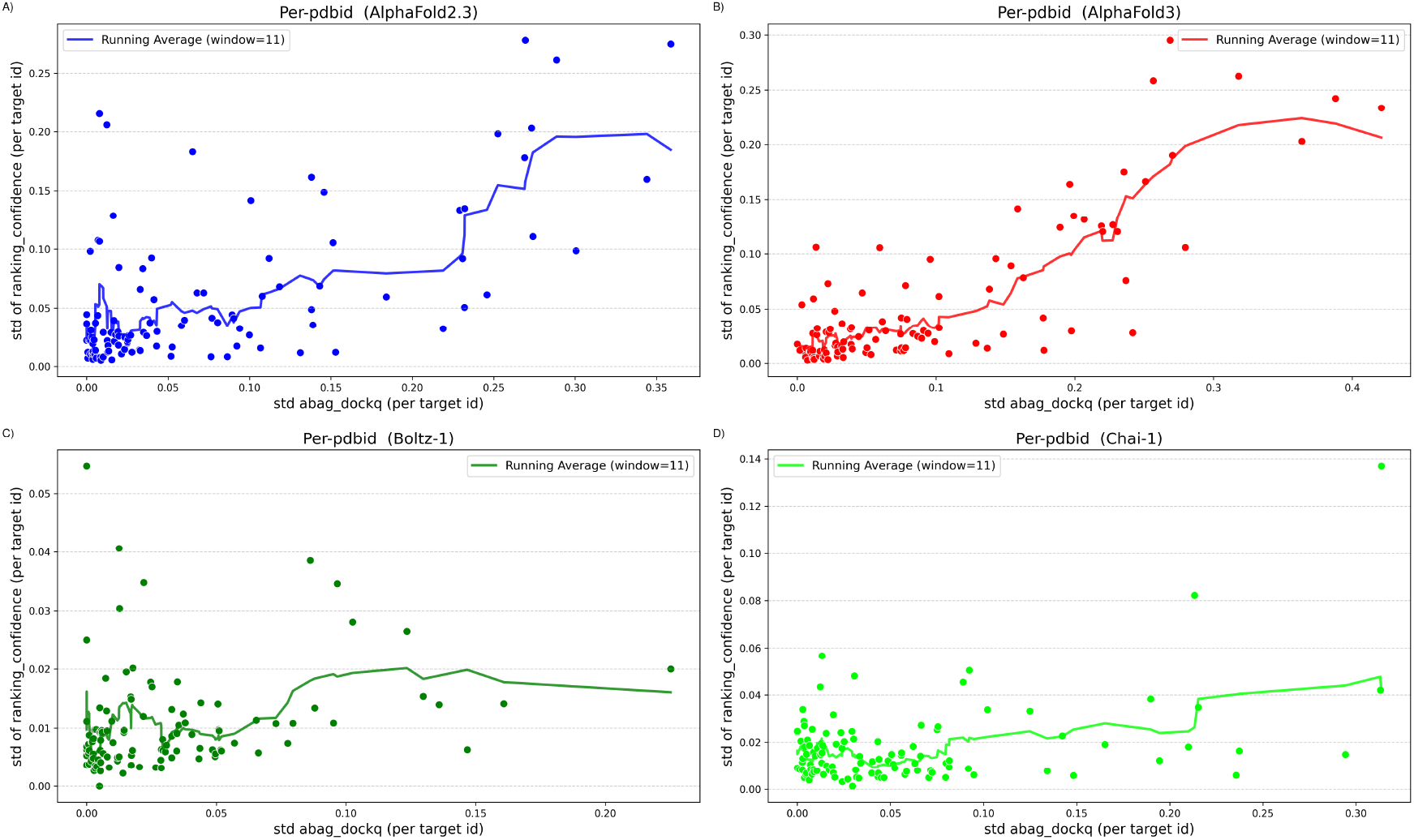
Each figure shows a scatter plot for the Standard Deviation of the ranking confidence scores versus the standard deviation of the DockQ for each target. A) AlphaFold2.3, B) AlphaFold3, C) Boltz-1, and D) Chai-1. The running average (over 11 samples) shows a minor trend that the variation is largest for “intermediately” easy targets.

**Fig. S6.**
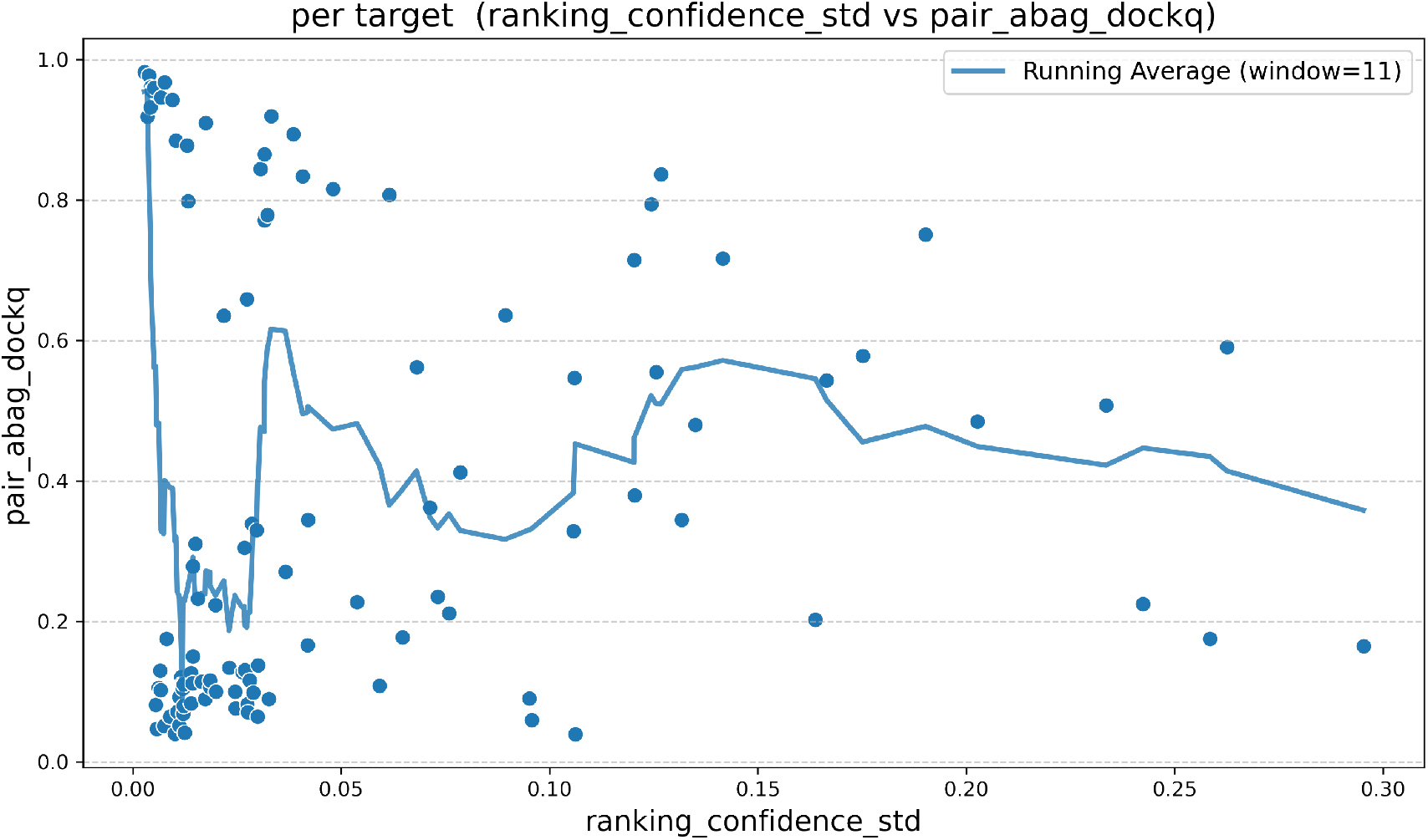
Variation of ranking confidence plotted against variation of the structures, as measured by average DockQ similarity between the generated models.

**Fig. S7.**
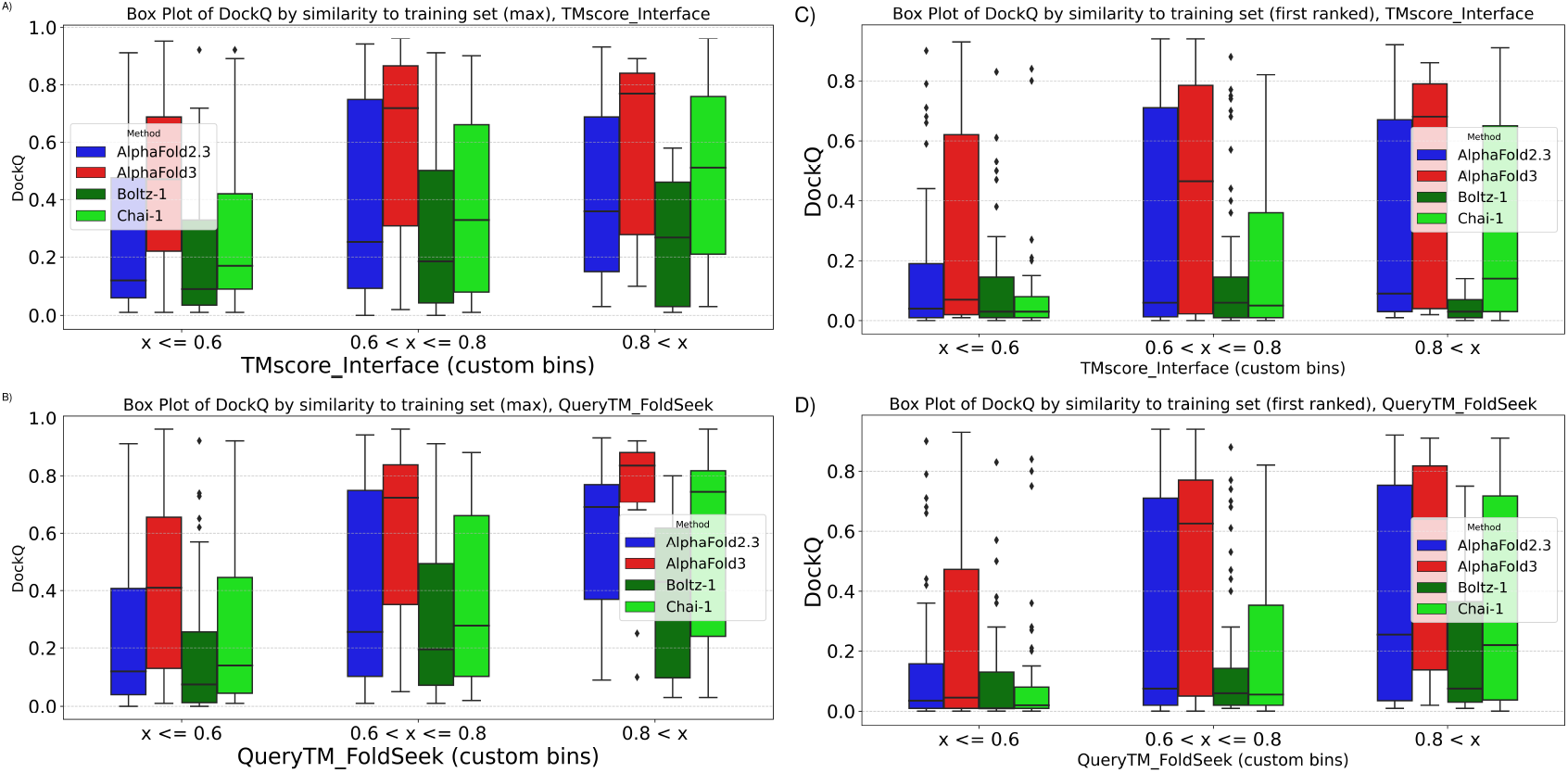
Prediction performance vs. similarity to the training data for AlphaFold2.3 and AlphaFold2 using the FoldSeek *query TMscore* (A,B) or *Interface TM-score (C,D)*. Plotted for the max score in A and C and for the first-ranked scores in B and D.

**Fig. S8.**
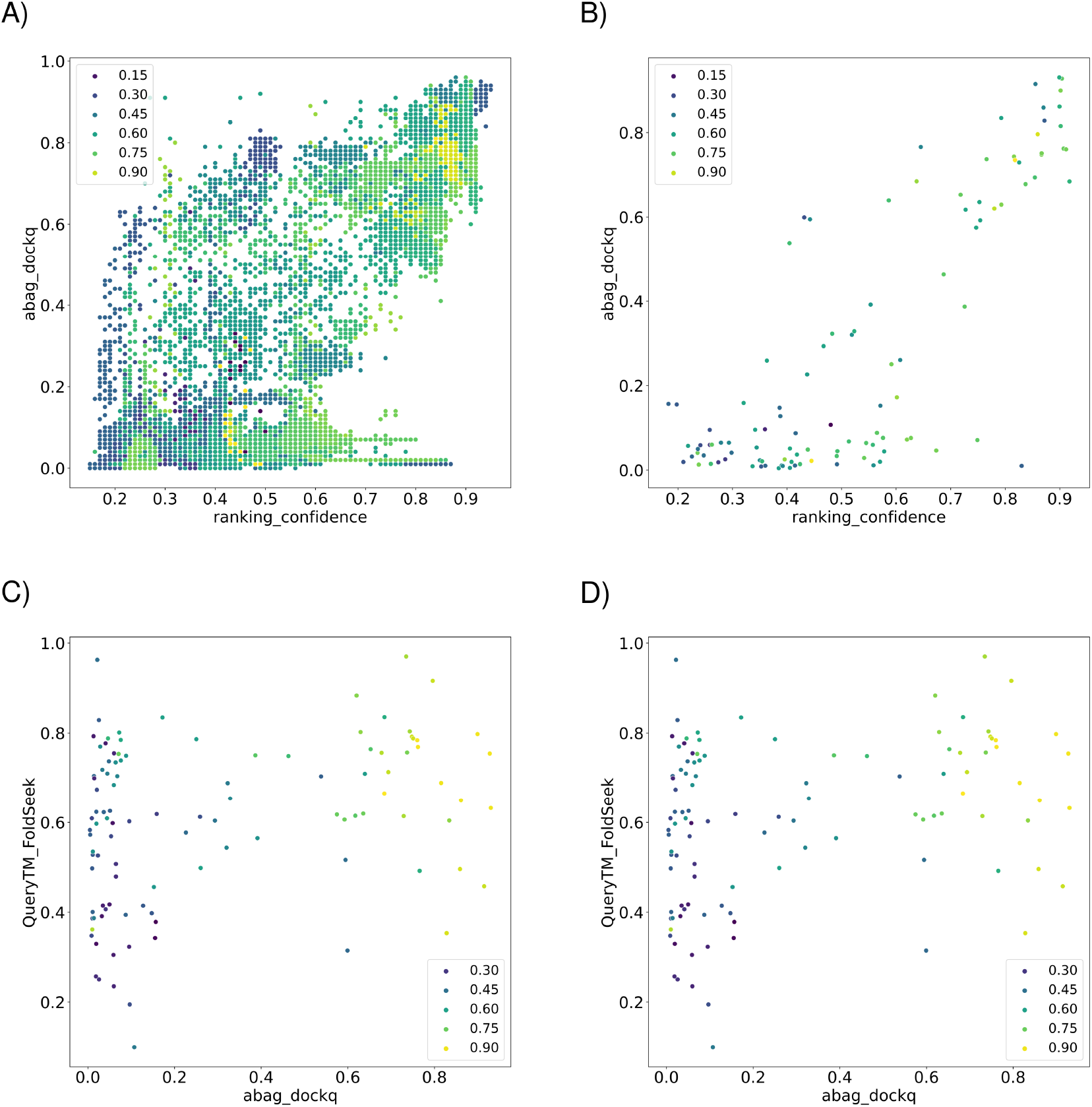
A) Ranking confidence vs DockQ for all models, colouring is by maximum similarity to the training set (TM-score). B-D) The same data plotted for the mean for each target, using different projections.

**Fig. S9.**
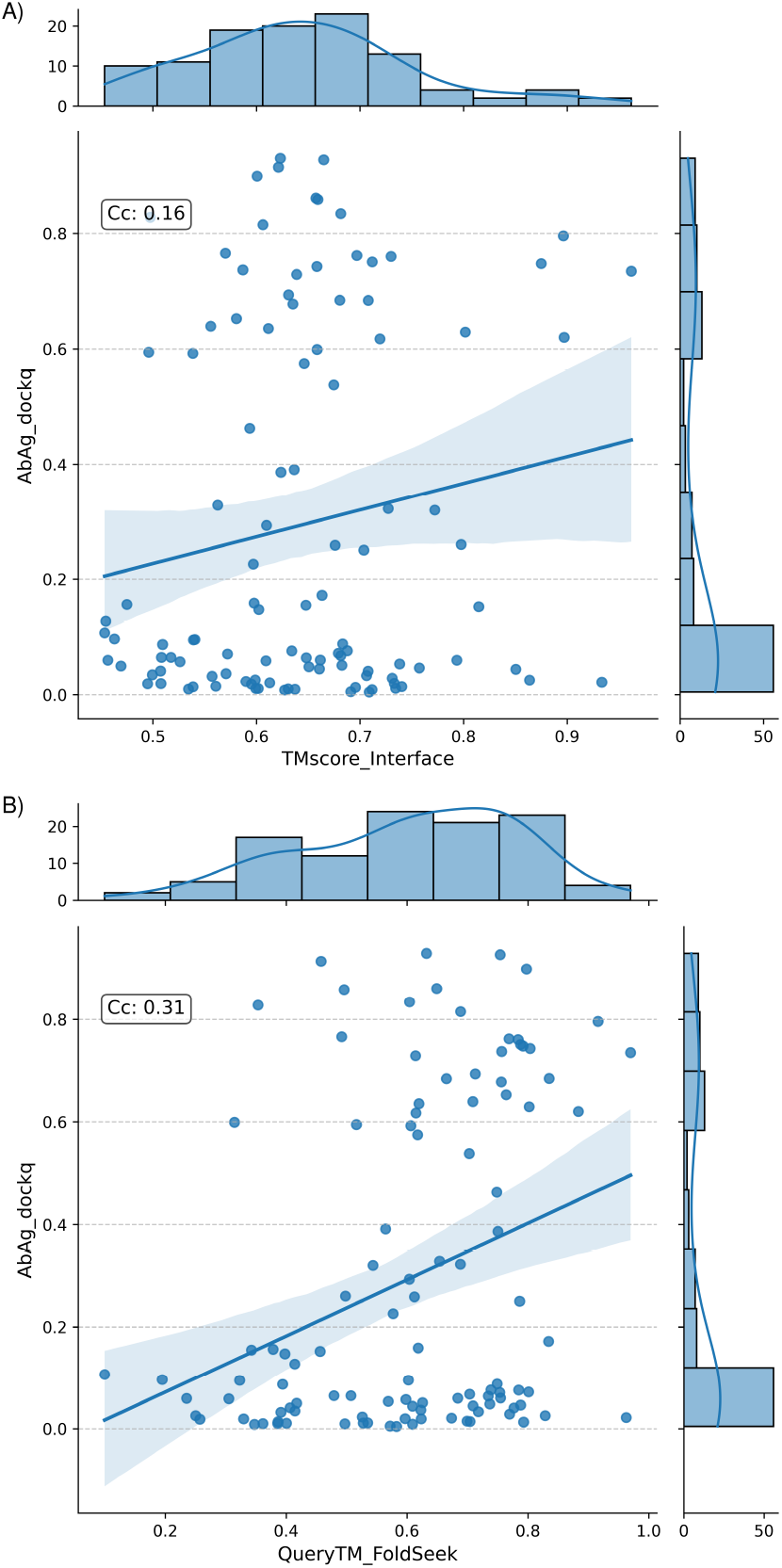
Scatter plot of mean DockQ scores vs similarity to training set using interface TM-score similarity (A) or Query TMscore similarity (B) for AlphaFold3.

**Fig. S10.**
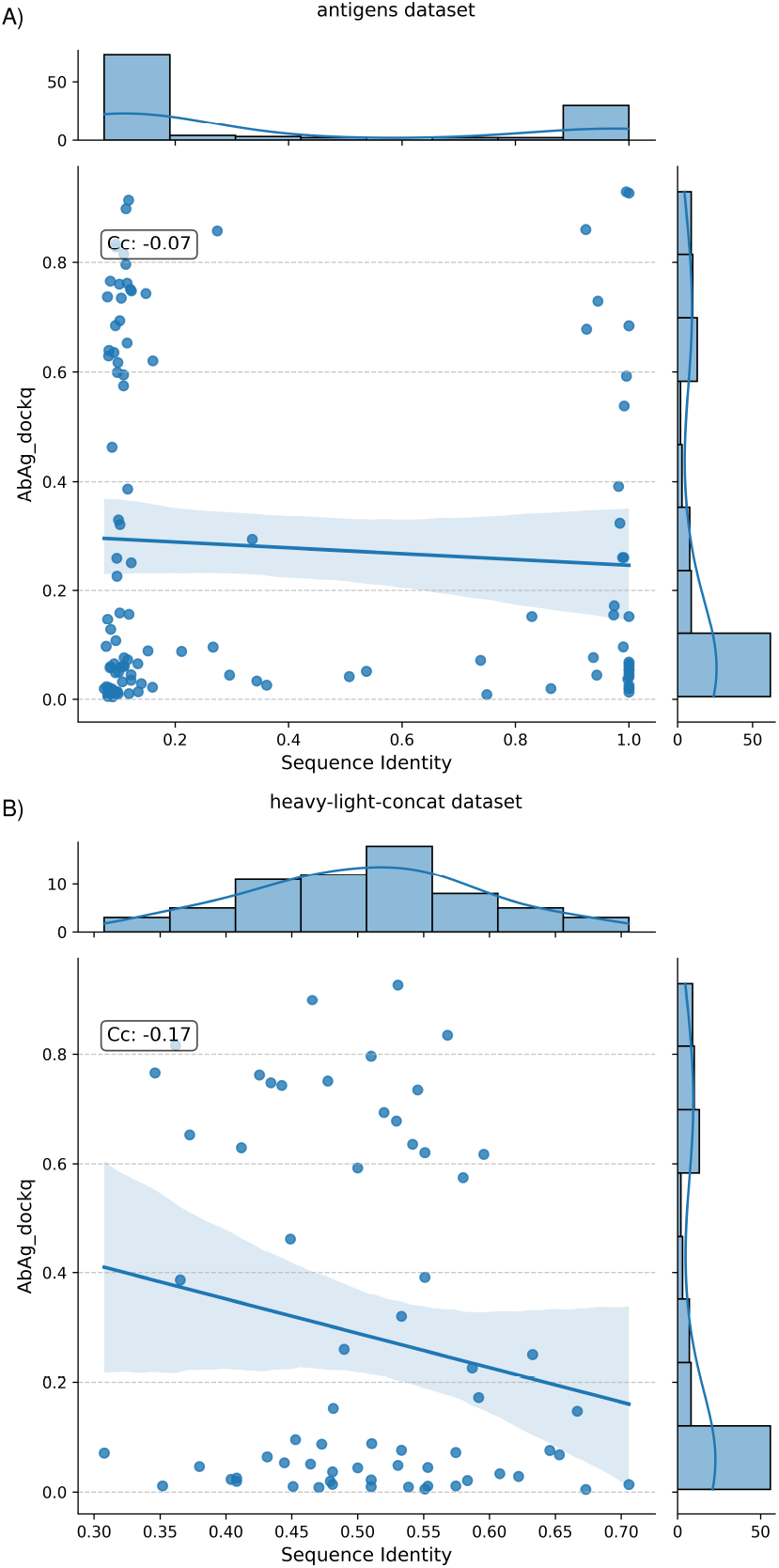
Scatter plot of mean AbAg-DockQ scores vs maximum sequence identity of (a) antigens and (B) concatenated CDR loops to the training set, for AlphaFold3.

**Fig. S11.**
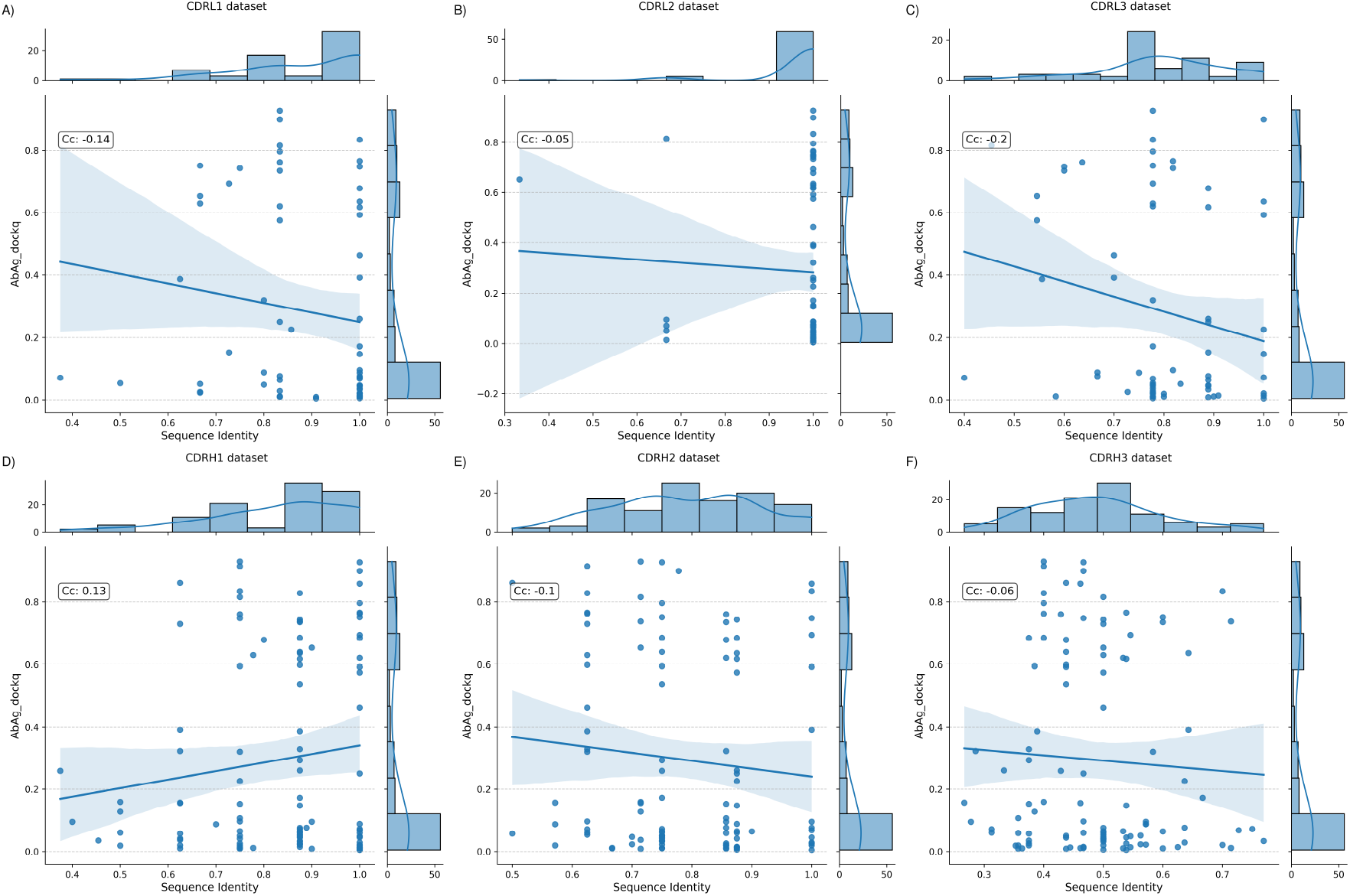
Scatter plot of mean AbAg-DockQ scores vs maximum sequence identity for individual loops to the training set, for AlphaFold3.

**Fig. S12.**
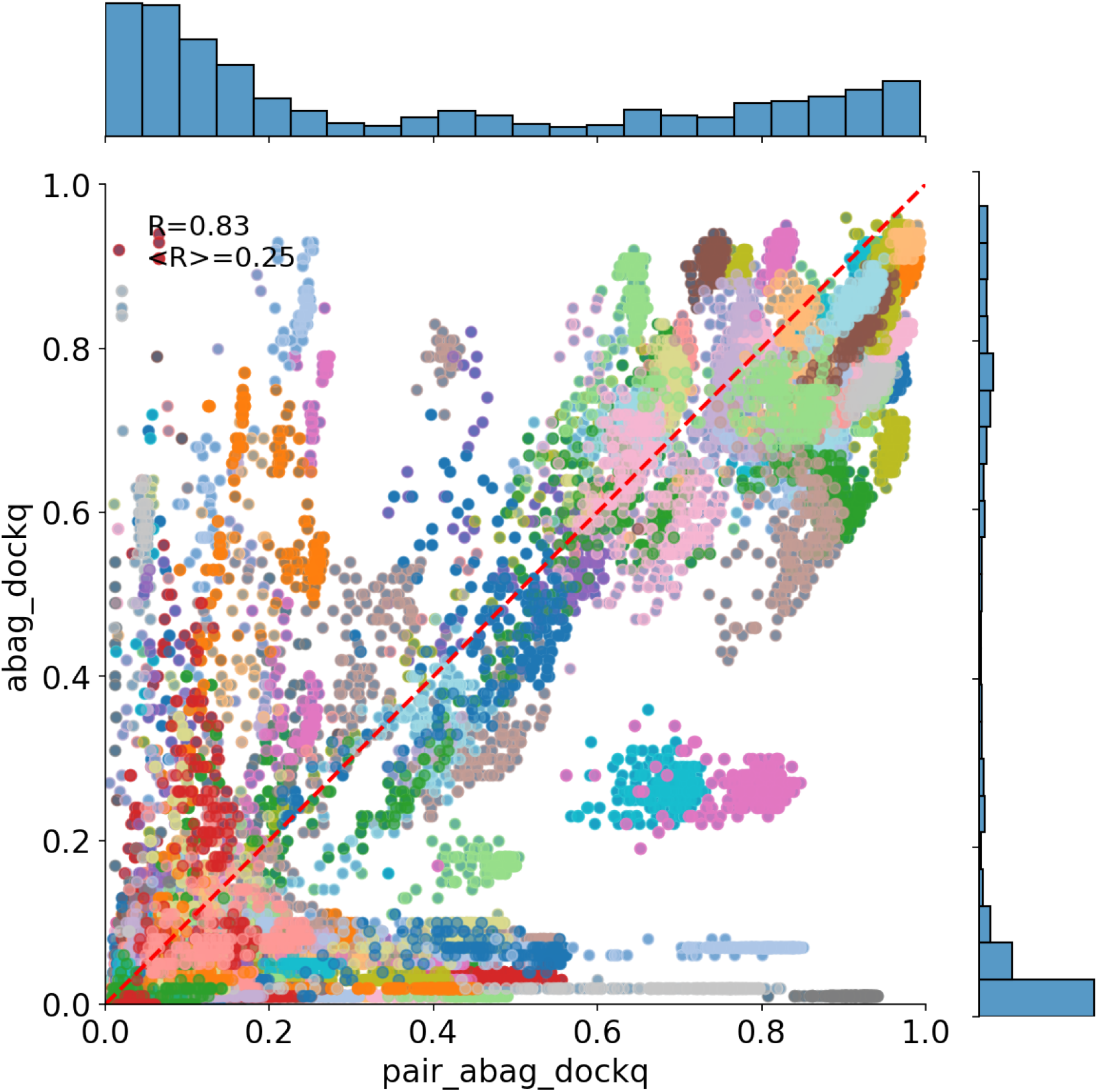
Scatterplot of paired DockQ vs DockQ for all models. Each target is colored separately. The correlation overall is 0.85, while the correlation per target is 0.19.

**Fig. S13.**
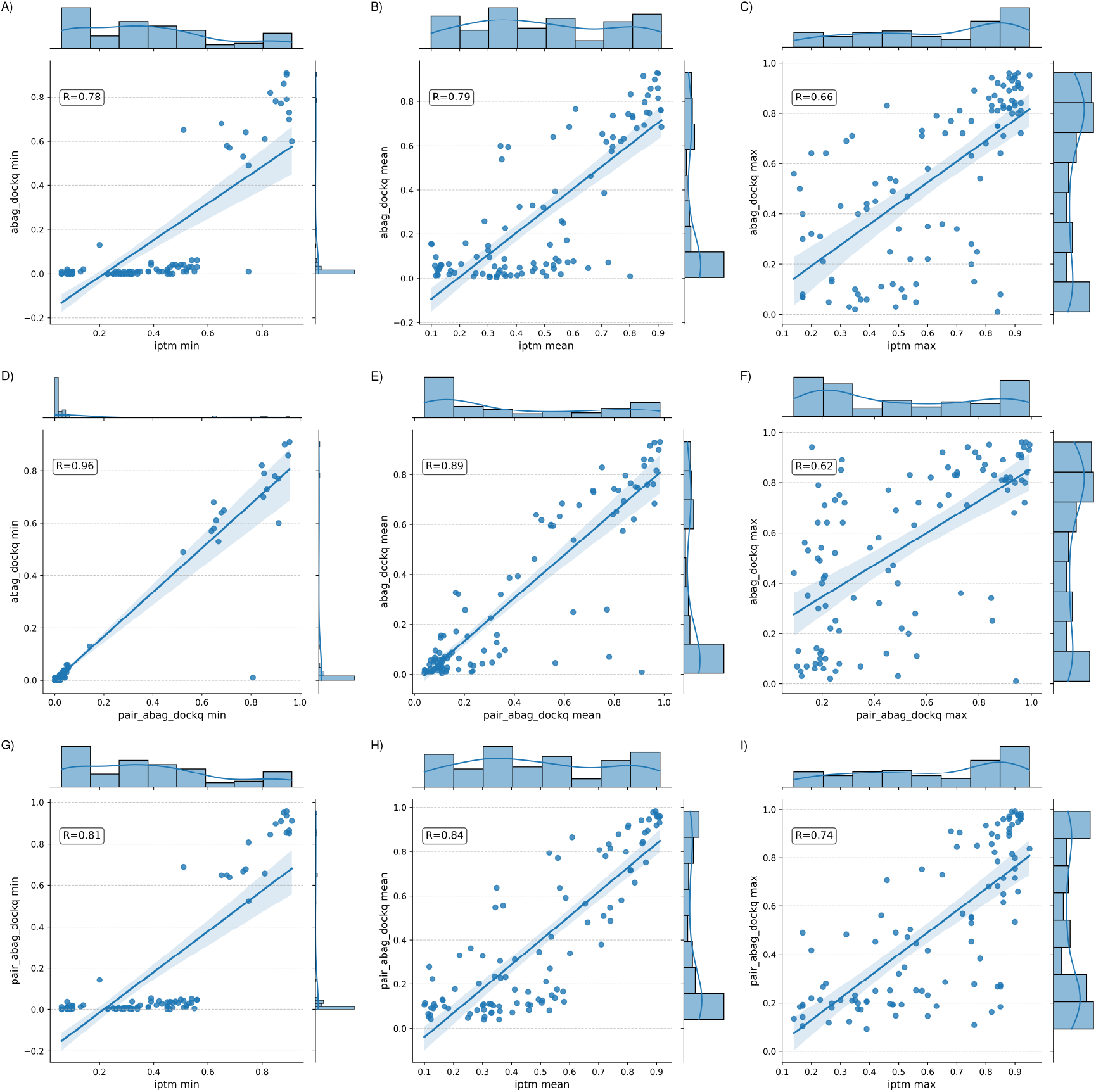
Comparison of DockQ, Paired DockQ and ipTM for min, mean, and max scores for each target. Overall, all three measures correlate well, particularly when using the min or mean estimates, but Paired DockQ is a better predictor of DockQ than ipTM. One can also note that there is one notable outlier (8us3) that has a high ipTM and Paired DockQ scores but a very low DockQ score, i.e. it is a consistent high-scoring prediction that is wrong, a false positive, no matter what method is used to estimate its quality. The reason 8u3s is wrong is because of its annotation in SabDAB, here the annotated unit corresponds to the assymetric unit, which do not correspond to the the biological assembly.

**Fig. S14.**
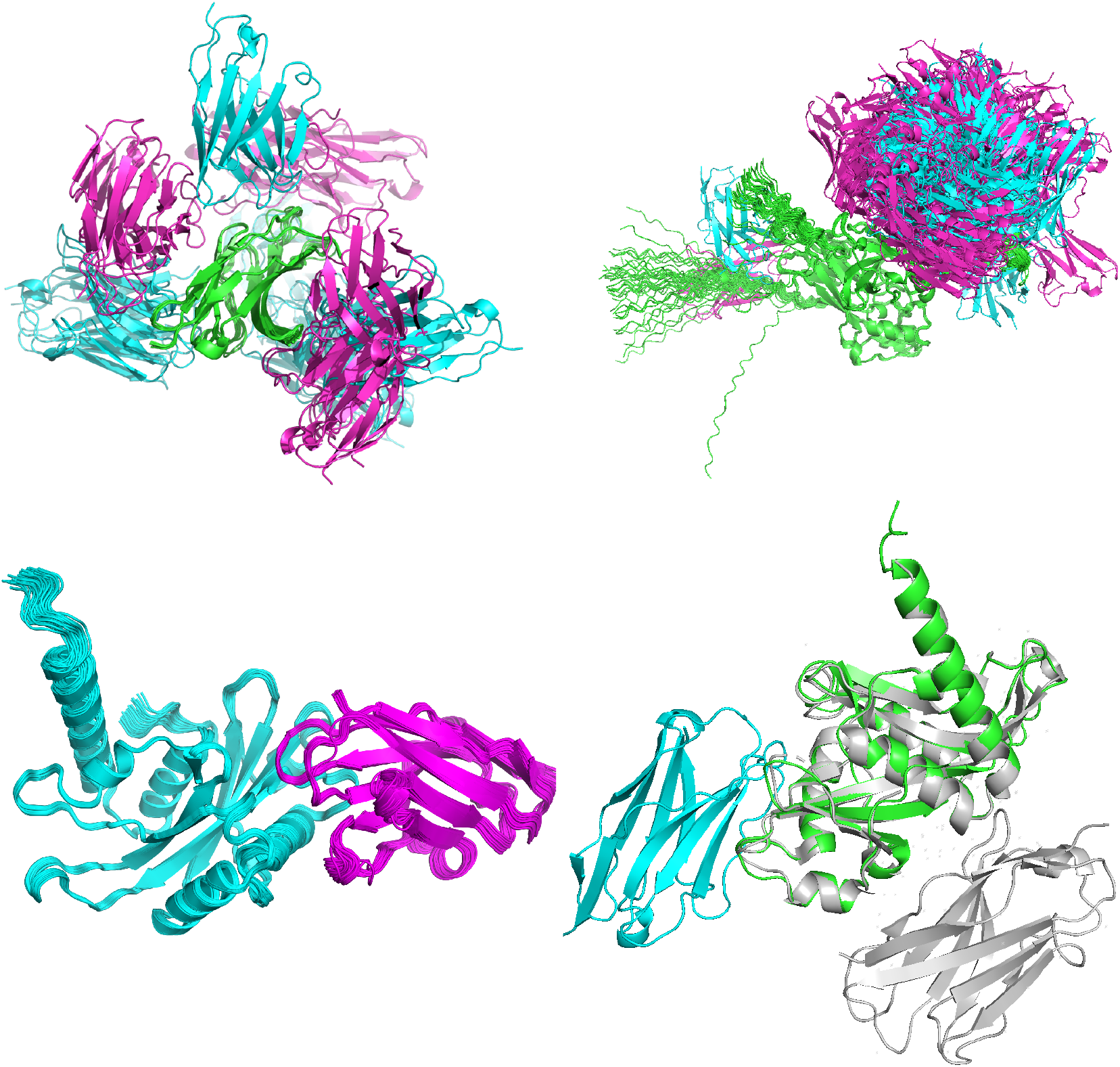
Example of predictions (all 40 sample 1 models) of predictions using AlphaFold3. All models are superposed on the antigen. A) 8hit (average Paired DockQ= 0.11, ipTM=0.54), B) 8jg5 (average Paired DockQ=0.09, ipTM=0.3 ) and C) 8U3S (average Paired DockQ= 0.91, ipTM=0.80) D) One prediction of 8U3S against the pdb file (in grey).

**Fig. S15.**
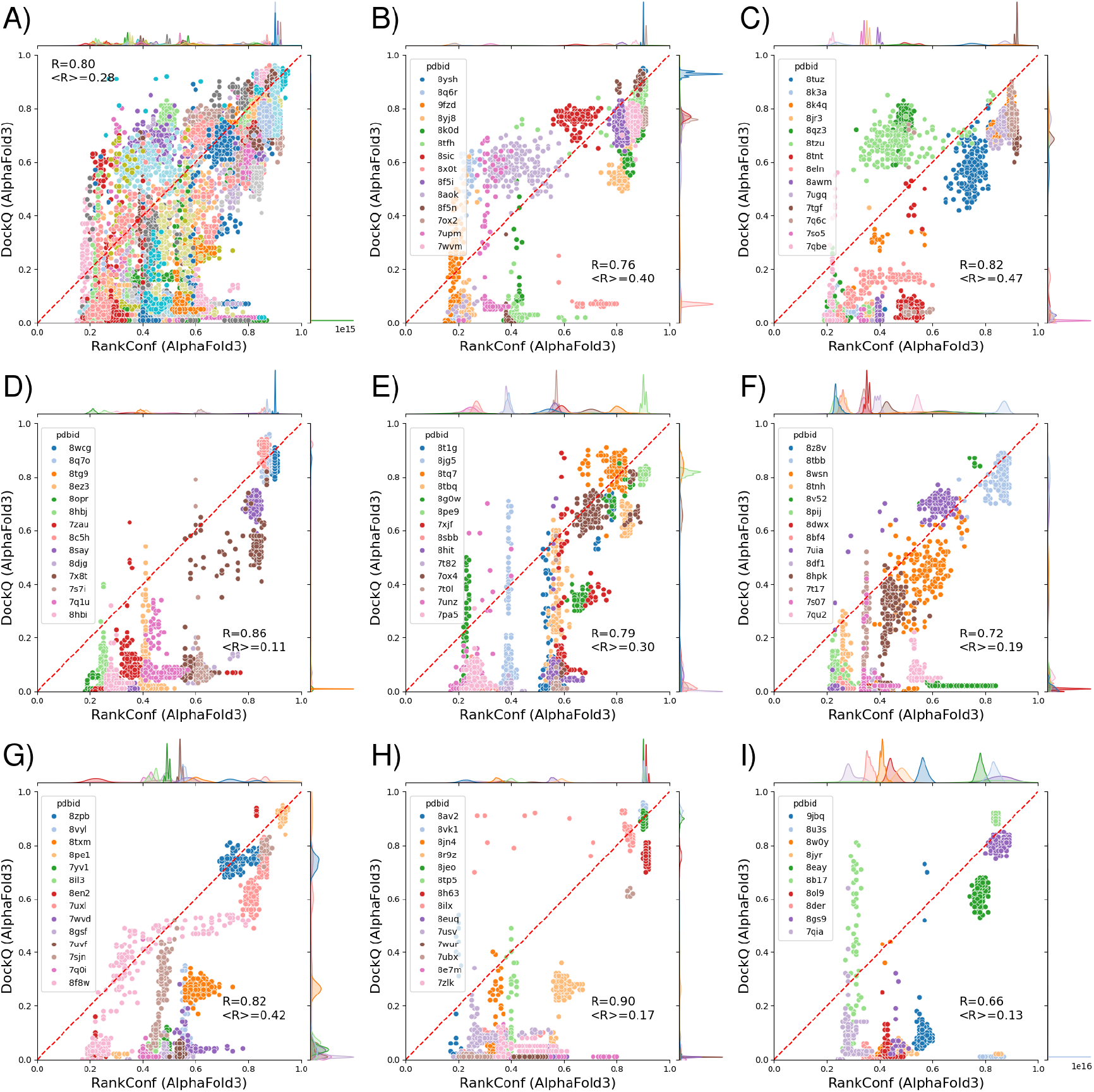
Scatter plot of Ranking Confidence vs AbAg-DockQ for individual targets. Ten targets per group, each coloured differently. In Figure A all are plotted together.

**Fig. S16.**
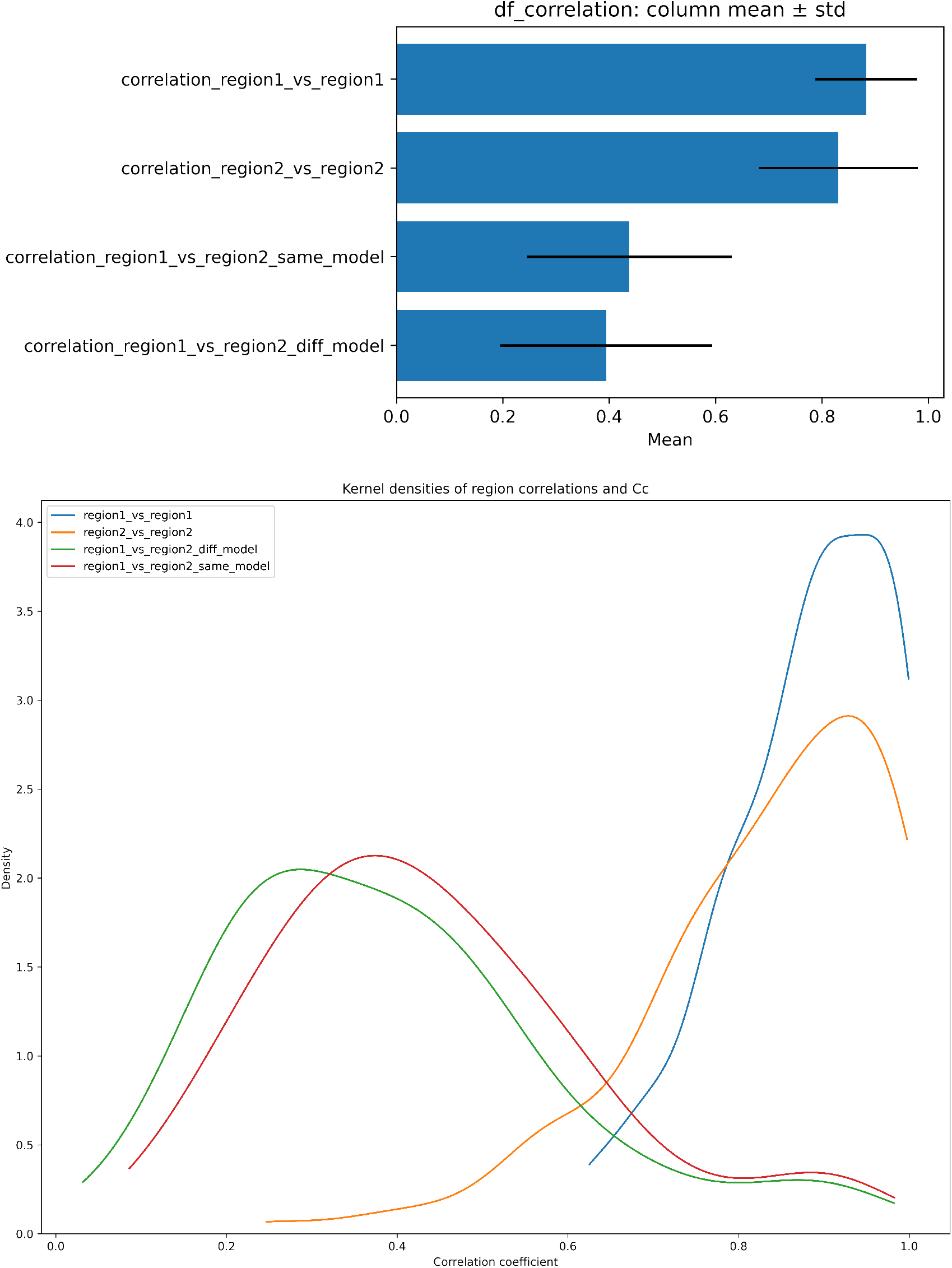
Top Figure: Correlation between the same and different regions for all models generated for one target. Region 1 is the upper region as shown in Figure 7, i.e. the PAEs when aligned on the antigen, while Region2 relates to the PAEs when aligned on antibody. Bottom Figure: The same distributions in a density plot.

**Fig. S17.**
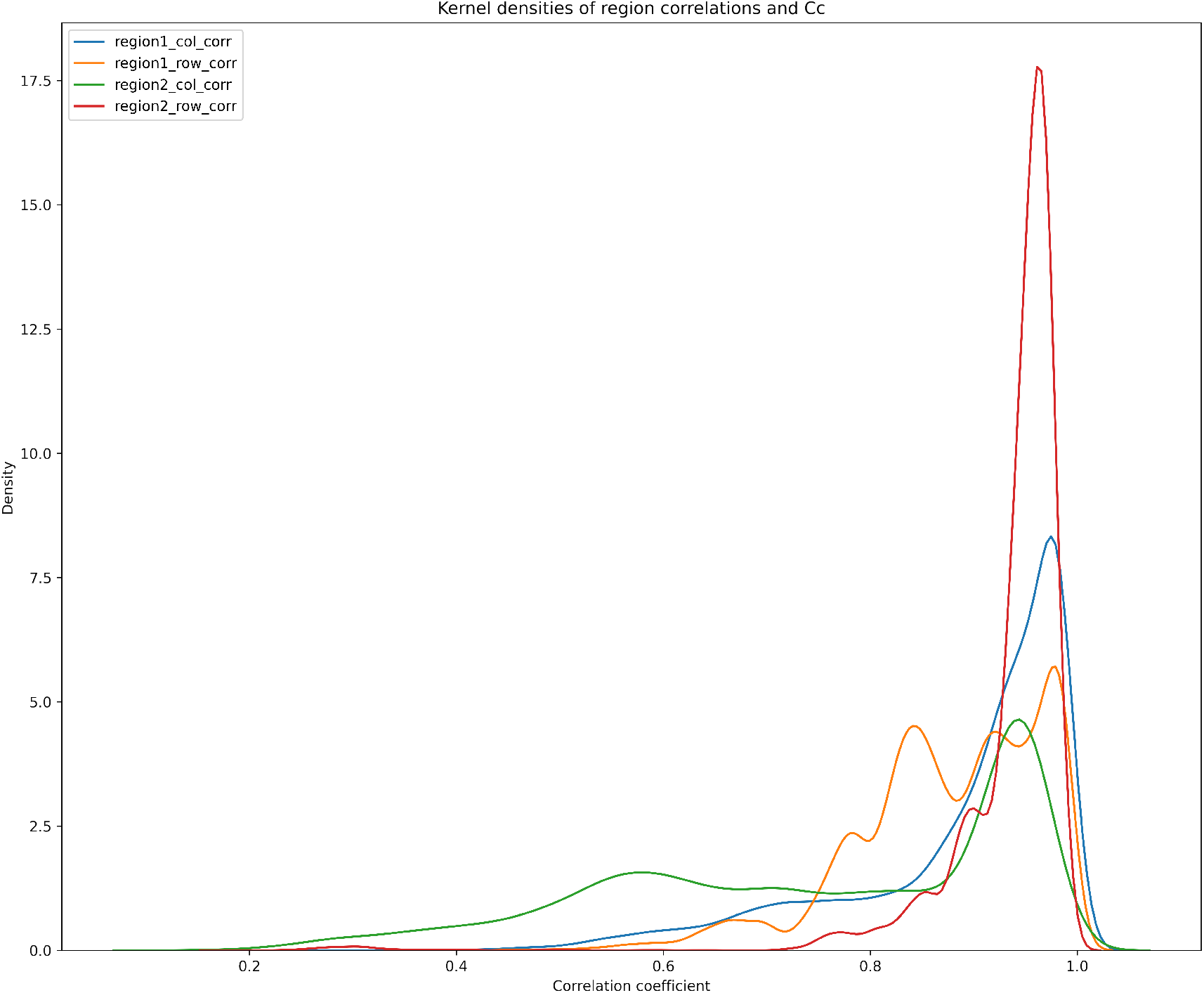
Correlation within rows/columns in region1 and region2, i.e. how similar are the PAEs when “aligning” on different residues in the antibody (region2) or antigen (region1). The highest correlations are found for region2 row correlation, followed by region1 column correlation, which are the PAEs for residues in the antigen; i.e., there are residues in the antigen that always have higher/lower PAEs than the others.

## Notes

https://zenodo.org/records/15764903

### Competing Interest Statement

The authors have declared no competing interest.

### Summary of Updates

Updated text and added supplementary material

## References

1. R.-M. Lu, et al., Journal of Biomedical Science 27(1), 1 (2020).

2. J. Jumper, et al., Nature 596(7873), 583 (2021).

3. P. Bryant, G. Pozzati, A. Elofsson, Nat Commun 13(1), 1265 (2022).

4. M. Saluri, M. Landreh, P. Bryant, PLoS Comput Biol 21(6), e1013168 (2025).

5. R. Yin, B. G. Pierce, Protein Sci 33(1), e4865 (2024).

6. B. Wallner, Bioinformatics 39(9) (2023).

7. N. Raouraoua, et al., Nat Comput Sci 4(11), 824 (2024).

8. Y. Kalakoti, B. Wallner, Commun Biol 8(1), 373 (2025).

9. D. Del Alamo, D. Sala, H. S. Mchaourab, J. Meiler, Elife 11 (2022).

10. R. Evans, et al., bioRxiv p. 2021.10.04.463034 (2022).

11. K. M. McCoy, M. E. Ackerman, G. Grigoryan, Protein Sci 33(9), e5127 (2024).

12. C. Schneider, M. I. J. Raybould, C. M. Deane, Nucleic Acids Res 50(D1), D1368 (2022).

13. M. Mirdita, et al., Nature Methods 19(6), 679 (2022).

14. S. Mukherjee, Y. Zhang, Nucleic Acids Res 37(11), e83 (2009).

15. U. Zhang, J. Sjolnick, Proteins pp. 702–710 (2004).

16. C. Mirabello, B. Wallner, Bioinformatics 40(10) (2024).

17. S. Basu, B. Wallner, PLoS One 11(8), e0161879 (2016).

18. R. L. Dunbrack, bioRxiv p. 2025.02.10.637595 (2025).

19. W. Zhu, A. Shenoy, P. Kundrotas, A. Elofsson, Bioinformatics 39(7), btad424 (2023).

20. M. van Kempen, et al., Nat Biotechnol 42(2), 243 (2024).

21. W. Kim, et al., Nat Methods (2025).

22. C. Zhang, M. Shine, A. M. Pyle, Y. Zhang, Nat Methods 19(9), 1109 (2022).

23. G. Pozzati, et al., Bioinformatics 38(4), 954 (2022).

24. J. Abramson, et al., Nature 636(8042), E4 (2024).

25. J. Wohlwend, et al., bioRxiv p. 2024.11.19.624167 (2025).

26. C. Discovery, et al., bioRxiv p. 2024.10.10.615955 (2024).

27. N. Raouraoua, M. F. Lensink, G. Brysbaert, Proteins (2025).

28. J. K. Varga, S. Ovchinnikov, O. Schueler-Furman, arXiv p. 2412.15970 (2024).

